# Metabolic breakdown of non-small cell lung cancers by mitochondrial HSPD1 targeting

**DOI:** 10.1101/2021.03.09.434573

**Authors:** Beatrice Parma, Vignesh Ramesh, Paradesi Naidu Gollavilli, Aarif Siddiqui, Luisa Pinna, Annemarie Schwab, Sabine Marschall, Shuman Zhang, Christian Pilarsky, Francesca Napoli, Marco Volante, Sophia Urbanczyk, Dirk Mielenz, Henrik Daa Schrøder, Marc Stemmler, Heiko Wurdak, Paolo Ceppi

## Abstract

The identification of novel targets is of paramount importance to develop more effective drugs and improve the treatment of non-small cell lung cancer (NSCLC), the leading cause of cancer-related deaths worldwide. Since cells alter their metabolic rewiring during tumorigenesis and along cancer progression, targeting key metabolic players and metabolism-associated proteins represents a valuable approach with a high therapeutic potential. Metabolic fitness relies on the functionality of heat shock proteins (HSPs), molecular chaperones that facilitate the correct folding of metabolism enzymes and their assembly in macromolecular structures. Here, we show HSPD1 (HSP60) as a survival gene ubiquitously expressed in NSCLC and associated with poor patients’ prognosis. HSPD1 knockdown or its chemical disruption by the small molecule KHS101 induces a drastic breakdown of oxidative phosphorylation, and suppresses cell proliferation both in vitro and in vivo. By combining drug profiling with transcriptomics and through a whole-genome CRISPR/Cas9 screen, we demonstrate that HSPD1-targeted anti-cancer effects are dependent on OXPHOS and validated molecular determinants of KHS101 sensitivity, in particular, the creatine-transporter SLC6A8 and the subunit of the cytochrome c oxidase complex COX5B. These results highlight mitochondrial metabolism as an attractive target and HSPD1 as a potential theranostic marker for developing therapies to combat NCSLC.

**Significance:** HSPD1 elimination or disruption interferes with NSCLC metabolic activity causing a strong OXPHOS-dependent energetic breakdown, which the cancer cells fail to overcome, highlighting HSPD1 as a potential theranostic marker for improving lung cancer therapy.

## INTRODUCTION

Lung cancer is the most commonly diagnosed cancer worldwide and the main cause of cancer-related death in both men and women (1). Non-small cell lung cancer (NSCLC) is lung most frequent type occurring in about 85% of cases (2), mainly presenting with an adenocarcinoma (LUAD) or squamous cell lung cancer (SqCC) histology (3). The standard of care for NSCLC has considerably changed in the last decade with the introduction of target therapies and immunotherapies. These, in synergy with chemo/radiotherapy, allowed to improve the clinical outcome of a fraction of NSCLC patients, in some cases providing a durable disease stabilization (4). Nevertheless, relapse and occurrence of treatment-resistant phenotypes are still very common, contributing to the current suboptimal scenario of 60 to 70% overall survival for early-stage and between 0 to 10% for patients carrying the advanced disease (5). This unsatisfactory picture strongly reinforces the need for developing new drugs targeting other fundamental lethal properties of LC cells.

It has long been established that cancer cells alter their metabolism to sustain their increased energetic demands and fuel the malignant phenotype (6). In contrast, or rather as an expansion of the so-called “Warburg effect”, it is now clear that cancers not only depend on glycolysis for their growth, but also rely on mitochondrial respiration (7), enabling a hybrid or “plastic” behavior in which an opportunistic energetic adaptability determines or facilitate survival and aggressiveness (8). However, the metabolic alterations frequently found in tumor cells could be turned into therapeutic opportunities, as the newly acquired cancer-specific vulnerabilities or dependencies have been found to represent, in some cases, excellent drug targets (9,10). Recent pivotal studies have shown the power of this approach to induce a significant tumor growth arrest (11-13).

Many key cellular metabolic processes are dependent on the functionality of heat shock proteins (HSPs) (14). HSPs are highly conserved ATP-dependent molecular chaperons mainly involved in maintaining the correct protein structure, for instance avoiding mis-folding or aggregation and protecting cell integrity under stressful conditions (15). High levels of HSPs have been found in different cancer cells, where they sustain higher metabolic demand and correct misfolded oncoproteins (16). The mitochondrial heat shock protein family D member 1 (HSPD1), also called HSP60, belongs to this family (17) and together with the co-chaperonin HSP10 plays an essential role in mitochondrial-imported proteins folding or refolding under mitochondrial stress (18). HSPD1 is involved in several diseases such as neurodegenerative disorders or cardiovascular diseases (19). In addition, HSPD1 carries a central role in cancer development, with either pro-survival or pro-apoptotic functions reported in different tumor types (20). In NSCLC, HSPD1 It has been proposed as a biomarker, but little is known about its role or the effects of its suppression (21,22).

Here, we investigated the tumor-promoting role of HSPD1 and explored if its targeting could interfere with the malignant metabolic mechanisms underlying NSCLC. Our results revealed HSPD1 as a ubiquitous lung cancer dependency gene, which maintains the metabolic fitness and promotes growth/survival, thus highlighting it as an attractive therapeutic target for NSCLC.

## METHODS

### Cell culture and chemicals

All cell lines were cultured in media supplemented with 10% FBS, 1% Pen/Strep and 1% L-Glutamine (all from Sigma) at 37 °C and 5% CO_2_ in a humidified incubator. H460, H1299, H520, SK-MES-1 and Calu-1 (from ATCC) were cultured in RMPI-1640 (Sigma). H23 and H838 (both from ATCC) cells were cultured in RMPI-1640, supplemented with 1 mM sodium pyruvate (Gibco). A549 (ATCC), BEN (DSMZ), murine Lewis lung carcinoma cell line LL2, Ladi2.1 and Ladi3.1 cells (derived from p53fl/fl-LSL KRASG12D/+ mouse NSCLC model) were cultured in DMEM (Sigma). Human cells were STR-profiled, used between passages 3 and 15, examined for mycoplasma regularly (detection kit from Invivogen). A549-Cas9 cell line was generated by lentiviral transduction with lentiCas9-Blast (Addgene #52962) and selection with 100 μg/mL of blasticidin (Sigma) for 3 days. KHS101 hydrochloride (4888) was purchased from Tocris; cisplatin (CAS 15663-27-1), Z-VAD-FKM (CAS 187389-52-2) and 2-Deoxy-D-Glucose (sc-202010) were purchased from Santa Cruz Biotechnology; Adenosine 5′-triphosphate (ATP) disodium salt hydrate (A1852-1VL), Necrostatin-1 (N9037-10NMG), Ferrostatin-1 (SML0583-5MG) and 1-Fluoro-2,4-dinitrobenzene (DNFB, D1529) were purchased from Sigma. HB072 was obtained as previously shown (23).

### Proliferation assay and in vitro drug treatment

For in vitro drug treatment, cells were plated in a 96-well plate in low density (5–10% initial confluence) and incubated overnight. On the next day, cells were treated either with vehicle or with the drugs. Plates were loaded in IncuCyte-Zoom (Essen Bioscience) and scanned every 2–4 hours. For each scan, phase contrast image was acquired from every well and was analyzed by IncuCyte Zoom software. IC_50_ value was calculated after 72 hours of treatment as expression of confluency percentage of treated cells normalized to vehicle control (100% confluency) and using Graphpad (nonlinear regression curve fitting model) to obtain the value. Proliferation assay was performed as described in (24). Growth curves were analyzed with 2-way ANOVA test.

### Cytotoxicity death assay

For cytotoxicity death assay, cells were seeded in 96-well plates in low density (5,000 cells/well) and incubated overnight. 1000X Cytotox Green Reagent or Caspase 3/7 Green Apoptosis Assay Reagent (Essen Bioscience) was diluted in medium and working dilutions of the drug were prepared in Cytotox Green or Caspase3/7 Green supplemented media. After treatment, plates were loaded in IncuCyte Zoom and images were acquired in real-time for phase to quantify growth. Activity of green reagent was simultaneously acquired at the green channel to quantify death. IncuCyte Zoom software was used for the analysis and data export

### Extracellular flux assays

Basal oxygen consumption rate (OCR) and basal extracellular acidification rate (ECAR) measurements were determined using the XFe96 Extracellular Flux Analyzer (Seahorse Bioscience/Agilent Technologies). Cells were seeded in specialized culture microplates at a density of 10,000-20,000 cells/well and prepared as describe in (25). For Mito Stress Test (Agilent kit 103015-100) 1 μM oligomycin, 1 μM FCCP, 1 μM rotenone and 1 μM antimycin were sequentially injected at regular intervals. OCR, indicator for mitochondrial respiration, and ECAR, indicative of glycolysis, were measured.

### CL-100 ProLiFiler screening

26 different lung cancer cell lines (**Supplementary Table 1**) were screened to determine their viability upon treatment with KHS101. The drug was added to the cells one day after seeding and the treatment was performed by nanodrop-dispensing using a Tecan Dispenser. 0.1% DMSO (solvent) and Staurosporine (1,0E-05 M) served as high control (100% viability) and low control (0% viability), respectively. After 72 hours of incubation with the compound, cell plates were equilibrated to RT for one hour and CellTiter-Glo^TM^ Luminescence Cell Viability Reagent (Promega) was added to the cell suspension. The luminescence was measured one hour later using a luminometer. IC_50_ values were express as percentage of proliferation in presence of solvent alone (100% = high control) as compared to cells treated with 1E-5M Staurosporine (0% = low control). IC_50_ calculation was performed using GraphPad Prism software with a variable slope sigmoidal response fitting model.

### Statistical analysis

Statistical tests were performed with the GraphPad software v.8 comparing groups of different conditions with replicates. In all tests, the statistical significance was set at p ≤ 0.05.

Additional methods are available in the Supplements.

## RESULTS

### HSPD1 is a fitness gene and a potential target for NSCLC

To understand whether HSPD1 could be an attractive target for NSCLC, we performed a fitness analysis from a previously published dataset of a pan-cancer and genome-wide CRISPR/Cas9 screening of 18,009 genes, which included n=21 adenocarcinoma (ADC) and n=11 squamous cell carcinoma (SqCC) cell lines (26). As a result, the HSPD1 gene scored very high, ranking 25^th^ in ADC and 71^th^ in SqCC cell lines, out of 805 and 559 total ‘fitness’ genes, respectively **(Fig. 1A)**. This initial observation was supported by the results from an independent dataset based on a large-scale RNAi screen of 7,837 genes performed in 52 NSCLC cell lines (27), in which HSPD1 resulted to be a significant cancer-dependent gene in over 70% of the cells **(Fig. 1B, Suppl. Table 2),** confirming the importance of this gene for NSCLC survival. This importance is also reflected in clinical significance. In fact, survival analysis on HSPD1 mRNA expression showed that the patients with high levels of HSPD1 have a worse prognosis compared to those with low HSPD1, in terms of overall **(Fig. 1C)**, relapse-free **(Fig. 1D)** and disease-specific survival **(Fig. 1E)**. We then investigated the prevalence of HSPD1 expression in tissue samples from a small cohort of NSCLC patients (n=20) by immunohistochemistry (IHC), and measured a positive expression in 100% of the cases **(Fig. 1F-I);** expression pattern was cytoplasmic granular or diffuse, with some degree of heterogeneity in terms of intensity (semi quantitatively scored in a three-levels scale), with very high protein levels identified in 45% of them **(Fig. 1J)**. In order to confirm HSPD1 expression in NSCLC cell lines grown *in vitro*, a panel of 10 different cell lines, including three mouse-derived (Ladi3.1, LL2 and Ladi2.1) (28), was subjected to western blot quantification, and the results indicated a detectable HSPD1 protein in all samples **(Fig.1K)**. Separated protein isolation from mitochondria and cytosolic fractions confirmed that HSPD1 is predominantly localized in the mitochondria in NSCLC cells **(Fig. 1L)**. Therefore, HSPD1 is a mitochondrial protein essential for NSCLC with a strong prognostic value and high prevalence of expression in tissue samples, making it a very valuable candidate for targeted inhibition.

**Figure 1.**
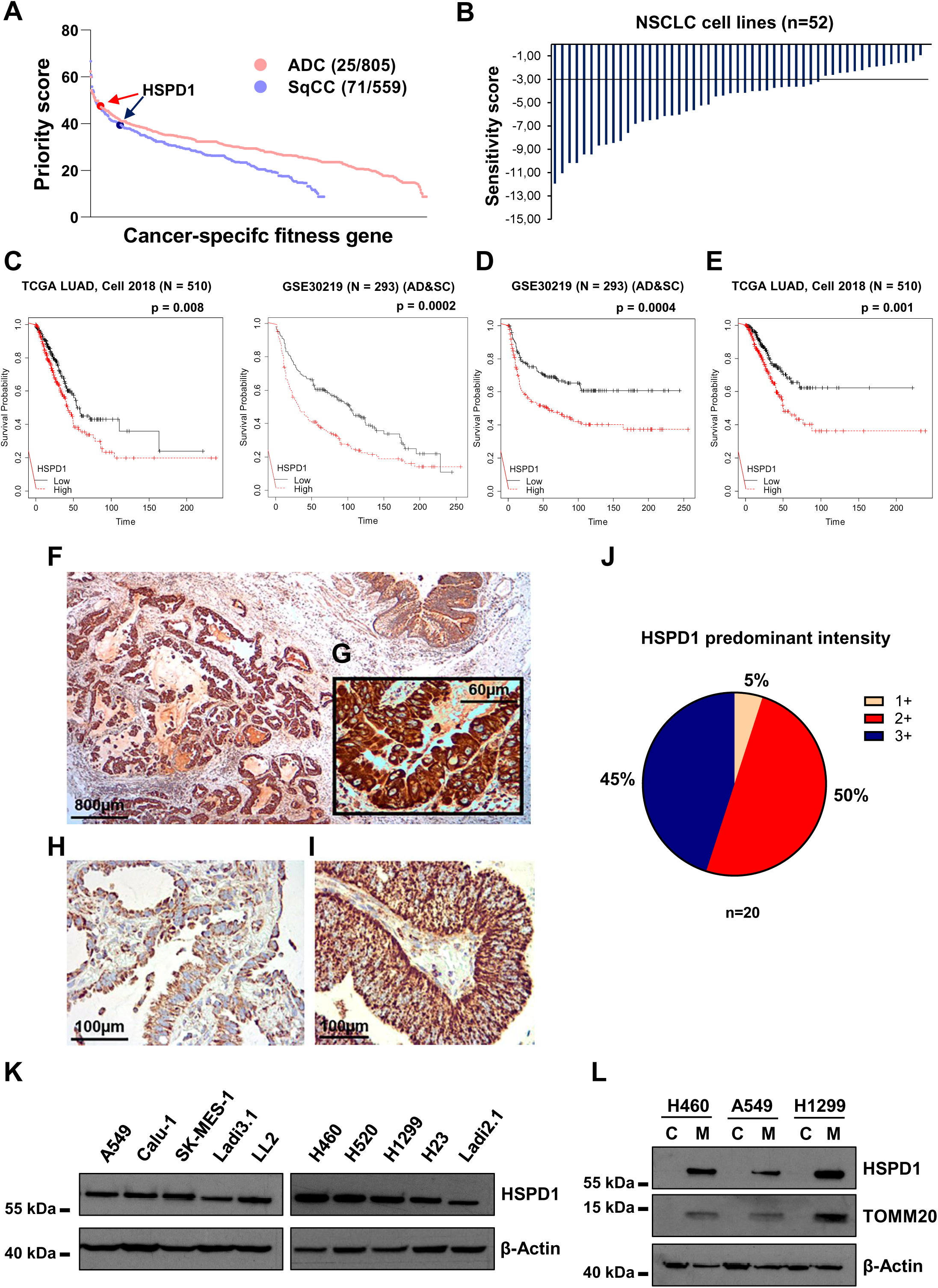
Heat shock protein HSPD1 is a fitness gene with prognostic power ubiquitously expressed in NSCLC tumors and cell lines. A) Fitness score for HSPD1 in lung adenocarcinoma (ADC) and squamous cell carcinoma (SqCC). Each dot represents a single gene. B) RSA (redundant siRNA activity) waterfall plot of HSPD1 sensitivity score from PROJECTDRIVE showing HSPD1 as an essential gene in non-small cell lung cancer (NSCLC) cell lines (n=52). RSA sensitivity score <-3. C) Overall survival analysis between the low- and high-HSPD1 groups in TCGA LUAD (N=510) and GSE30219 (N=293). P-value was calculated using log-rank test. Relapse-free (D) and disease-specific (E) survival analysis between low- and high-HSPD1 groups in GSE30219 (N=293) and TCGA LUAD (N=510), respectively. P-value was calculated using log-rank test. Time is expressed as months. IHC staining of HSPD1 in cancer tissues (F, G, H and I) derived from consecutive 20 lung cancer patients (peritumoral bronchial structure in upper right corner of panel F). J) Pie chart of HSPD1 predominant intensity in patient-derived adenocarcinoma samples. K) Western blot quantification of HSPD1 in a panel of different human (A549, Calu-1, SK-MES-1, H460, H520, H1299 and H23) and mouse (Ladi2.1, LL2 and Ladi3.1) NSCLC cells. β-Actin was used as loading control. L) Western blot analysis of HSPD1 in cytosolic (named as C) or mitochondrial (named as M) fraction. β-Actin was used as loading control and TOMM20 was used as control for cytosolic/mitochondrial fractionation.

### HSPD1 promotes NSCLC growth *in vitro* and *in vivo*

To study the function of HSPD1 in NSCLC cells *in vitro*, we analyzed the effects of its knockdown by transducing 5 different NSCLC cell lines with up to 5 independent shRNA sequences (called #44, #45, #46, #47 and #48) **(Fig. 2A, Suppl. Fig. 1A)** and evaluated proliferation through a real-time cell confluency assay (Incucyte system). Shortly after shRNA-based inhibition of HSPD1 expression, the cells showed a decrease in proliferation compared to the controls transduced with a scrambled/non-targeting shRNA, referred to as pLKO **(Fig. 2B and Suppl. Fig 1B)**. In particular, the phenotype was stronger with shRNA sequences providing higher level of HSPD1 suppression (like the #48). HSPD1 knockdown cells were not able to form colonies compared to the control cells when plated at lower density as observed over two weeks **(Fig. 2C-D and Suppl. Fig 1C-D)**. In line with a strong and durable effect on cell proliferation, a cell cycle analysis showed a significant increase in the percentage of cells in sub-G1 and G1 phases in HSPD1 knockdown cells, in contrast to a reduction in cells residing in S or G2-M phases **(Fig. 2E-H, Suppl. Fig. 1E-F)**.

**Figure 2.**
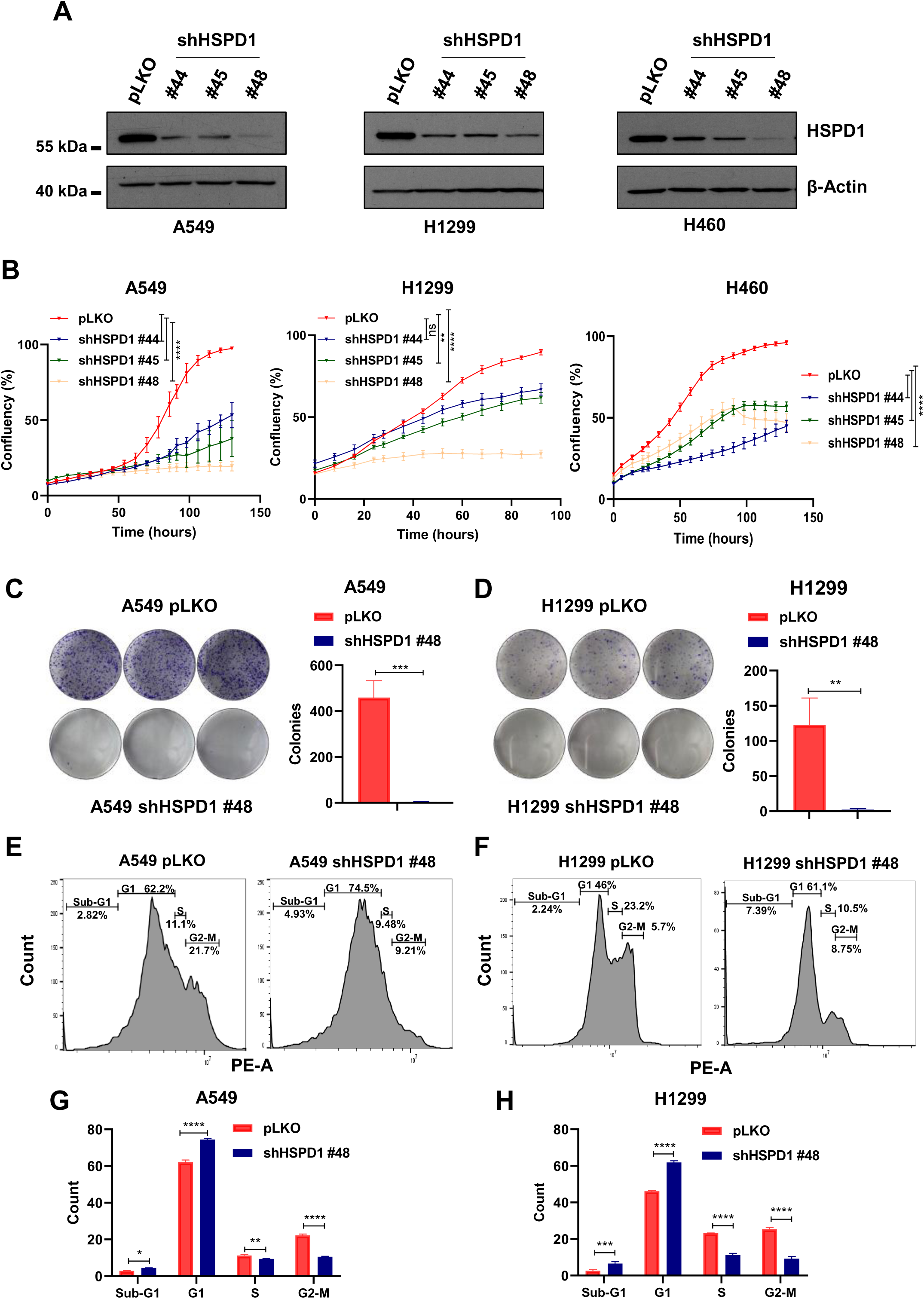
HSPD1 knockdown blocks cell proliferation and clonogenic ability of NSCLC cells. A) Western blot analysis of HSPD1 protein levels in A549, H1299 and H460 cells upon infection with 3 independent shRNA (#44, #45 and #48) targeting HSPD1 compared to scramble-infected cells (pLKO). β-Actin was used as loading control. B) Real-time proliferation curves of A549, H1299 and H460 infected with non-targeting pLKO or shHSPD1. Plotted is the cells’ confluency over time. P-values are from two-way ANOVA. Points (n=5) are average ± SD. *<0.05, **<0.01, ***<0.001, ****<0.0001. Colony formation of A549 (C) and H1299 (D) cells with pLKO or shHSPD1, stained with crystal-violet and quantified in triplicates. Scale bars are average ± SD. P-values are from unpaired *t*-test. **<0.01, ***<0.001. FACS plots of A549 (E) and H1299 (F) cells infected with pLKO or shHSPD1 and stained with PI for cell cycle analysis. Bar graph showing the % of cells in each cell cycle phase of A549 (G) and H1299 (H) cells upon infection with pLKO or shHSPD1. P-values are from two-way ANOVA. Scale bars (n=3) are average ± SD. **<0.01, ***<0.001.

HSPD1 has a critical function in mitochondrial metabolism (23). Therefore, in order to investigate whether the growth suppression observed with HSPD1 knockdown was linked to metabolic alterations, an extracellular metabolic flux analysis was performed using two different cell lines. Throughout the experiments, knockdown cells showed a significant reduction in their basal respiration and in their ATP-linked respiration, suggesting that the alterations in the mitochondrial metabolism may be responsible for the reduction in cell proliferation **(Fig. 3A-B)**. Finally, we tested if HSPD1 knockdown inhibited cell growth *in vivo.* To this end, we injected shHSPD1-expressing A549 and H1299 cells subcutaneously in NSG mice to evaluate the tumor growth rate over time. The results clearly indicated that HSPD1 knockdown cells formed very small and slower-growing tumors compared to control pLKO cells, in terms of both tumor volume and tumor weight **(Fig. 3C-H)**, an effect which was maintained for the entire duration of the experiment, confirming that HSPD1 is required for NSCLC growth.

**Figure 3.**
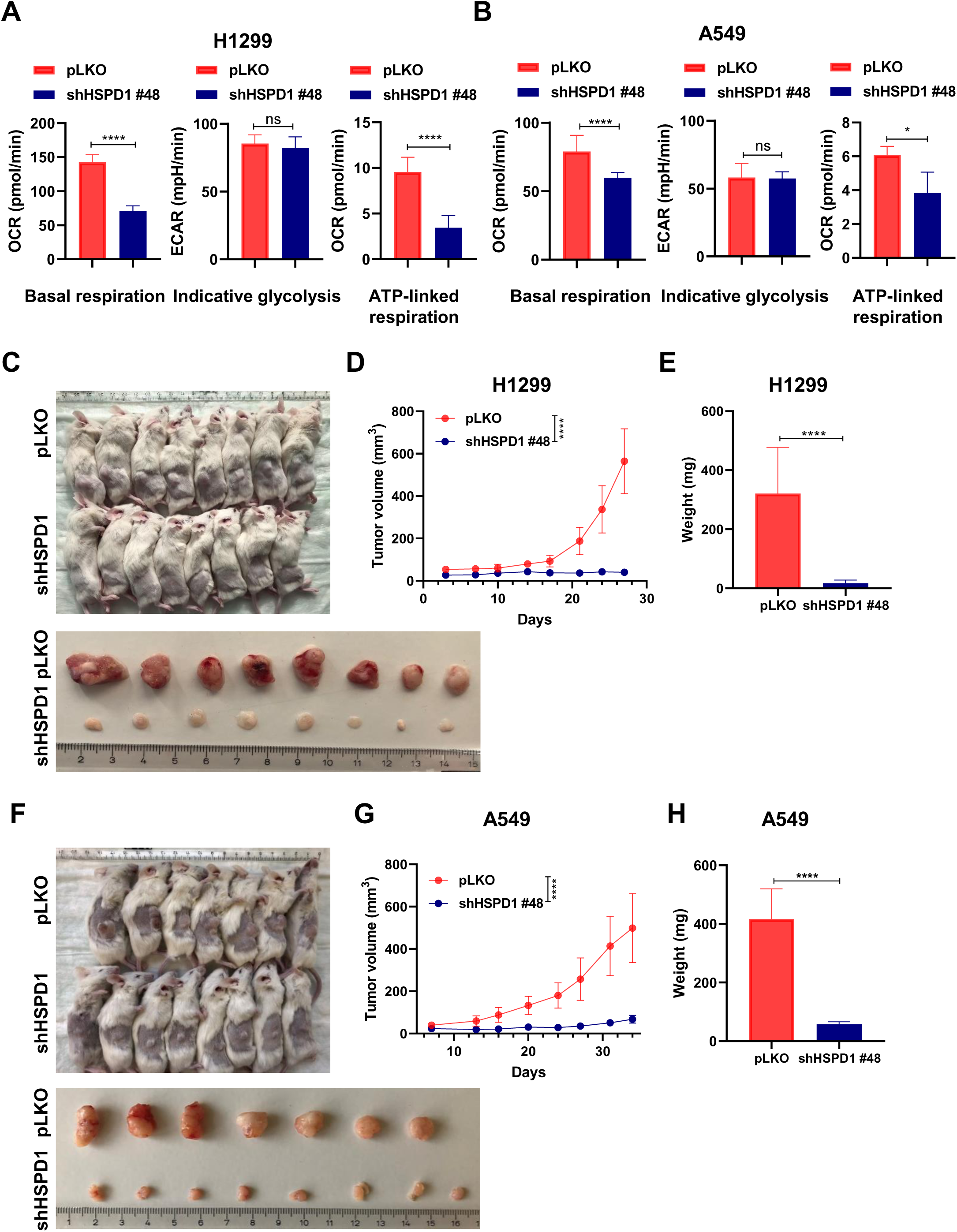
HSPD1 knockdown induces a metabolic breakdown and reduces tumor growth in vivo. Quantification of basal respiration, indicative glycolysis and ATP-linked respiration of H1299 (A) and A549 (B) shHSPD1 cells compared to pLKO cells. Bars (n=5) are average ± SD. P-values are from unpaired *t*-test. *<0.05, ****<0.0001. Images of tumors from NSG mice injected subcutaneously with H1299 (C) or A549 (F) pLKO or shHSPD1 cells. Graph showing tumor growth of H1299 (D) or A549 (G) shHSPD1 cells compared to pLKO cells. P-value is from two-way ANOVA. Points (n=8) are average ± SD. ****<0.0001. Bar graph showing weight of H1299 (E) and A549 (H) shHSPD1 tumors compared to pLKO tumors. Bars (n=8) are average ± SD. P-value is from unpaired *t*-test. ****<0.0001.

### The HSPD1-targeting small molecule KHS101 arrests NSCLC growth

KHS101 is a synthetic small molecule reported to impair the growth of glioblastoma cells *in vitro* and *in vivo* via inhibition of HSPD1 chaperone activity (23). As KHS101 leads to metabolic exhaustion and apoptosis in glioblastoma cells, we first tested as to whether the compound elicits cytotoxicity in NSCLC cell lines. After treatment with KHS101, 6 human NSCLC cell lines showed a significant reduction in cell proliferation that was dependent on the compound concentration **(Fig. 4A, Suppl. Fig. 2A-B)**. To verify the specificity of HSPD1 targeting, we exposed the cells to HB072, an HSPD1 non-targeting chemical analogue (23) **(Fig. 4B)**, which did not significantly affect cell growth **(Fig. 4C, Suppl. Fig. 3A)**. After 5 days of continuous drug treatment, NSCLC cells lost their clonogenic ability, as shown in a colony formation assay **(Fig. 4D-E, Suppl. Fig. 3B)**. In line with *HSPD1* knockdown experiments, cell cycle analysis showed a significant reduction in the percentage of cells in S and G2-M phases concomitant with an increase of cells in G1 phase **(Fig. 4F-I)**, as assessed 24 hours after KHS101 treatment. Finally, since many conventional anti-cancer therapeutic strategies fail to target non-proliferating cancer cells, we sought to test whether the KHS101-induced growth inhibitory effect was maintained altering the proliferative state of the cells in a controlled fashion. Therefore, after being synchronized overnight using a low FBS concentration (0.5%), the cells were treated with KHS101 in presence of media with increasing FBS, ranging from 0% to 5%. As a result, no differences were measured in KHS101 efficacy in the different conditions by a real-time proliferation assay **(Suppl. Fig. 3C)**, indicating that KHS101 is effective independently of cells proliferative status. In conclusion, these data indicate that chemical targeting of HSPD1 markedly reduces the growth of NSCLC cell lines.

**Figure 4.**
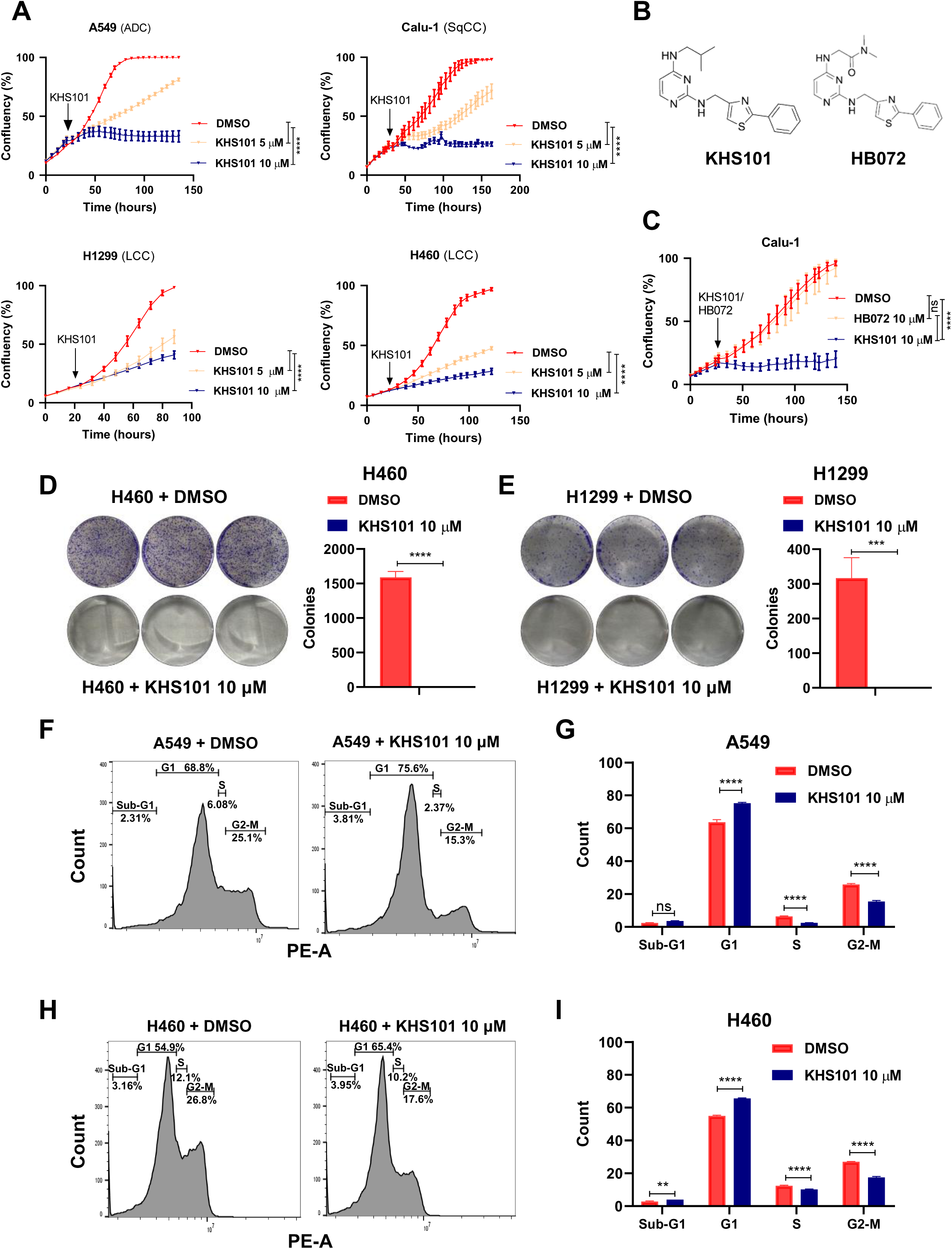
The small HSPD1-targeting molecule KSH101 induces cell growth arrest. A) Real-time proliferation curves of A549, Calu-1, H1299 and H460 treated either with vehicle (DMSO) or KHS101. P-values are from two-way ANOVA. Points (n=5) are average ± SD. ****<0.0001. B) Images of KHS101 and HB072 synthetic molecular structures. C) Real-time proliferation curves of Calu-1 cells treated either with KHS101 or with the corresponding inactive KHS101 analog (HB072) compared to vehicle-treated cells. P-values are from two-way ANOVA. Points (n=3) are average ± SD. ****<0.0001. Colony formation of H460 (D) and H1299 (E) treated for 5 days with KHS101 or vehicle and then left until forming visible colonies in drug-free media, then stained with crystal-violet and quantified in triplicates. Scale bars are average ± SD. P-values are from unpaired *t*-test. ***<0.001, ****<0.0001. FACS plots of A549 (F) and H460 (H) treated 24 hours with KHS101 and stained with PI for cell cycle analysis. Bar graph showing the % of cells in each cell cycle phase of A549 (G) and H460 (I) cells treated 24 hours with vehicle or KHS101. P-values are from two-way ANOVA. Scale bars (n=3) are average ± SD. **<0.01, ****<0.0001.

### KHS101 induces metabolic disruption in NSCLC cells leading to cell death

To test whether KHS101 affects energy metabolism in NSCLC cell lines, we performed an extracellular flux analysis 24 hours after KHS101 treatment. The results indicated a strong reduction in cellular basal respiration and in ATP-coupled respiration capacity. Concomitantly, a significant increase was observed for basal glycolysis **(Fig. 5A-B)**. We next asked as to how fast KHS101 would induce such alterations and treated the cells for shorter periods (ranging from 0.5h to 6h), observing that glycolytic activity increases simultaneously with OXPHOS reduction from less than one hour after treatment **(Suppl. Fig. 4A-B)**, and suggesting that the metabolic rewiring induced by KHS101 is an early event, as shown earlier (23). In addition to metabolic flux, cell morphology was found altered in cells exposed to 10 µM KHS101. Cells exposed to KHS101 presented an altered ultrastructure. After 96 hours of treatment severe swelling of mitochondria were seen with sparse presence of cristae. As sign of cellular degeneration, an increased number of secondary lysosomes were found. Moreover, there was a loss of microvilli **(Fig 5C)**. However, already after 48 hours mitochondria swelling was present, as well as loss of microvilli and in many cells increased number of lysosomes were present, too **(Suppl. Fig 4C)**. As a result of these alteration, and differently to HPSD1 knockdown, KHS101 treatment induced cell death, as indicated by the incorporation of Cytotox fluorescent green dye **(Fig 5D, Suppl. Fig. 4D)**. To test if the affected cells were undergoing apoptosis, we performed a caspase 3/7 activity assay, which showed specific activation of apoptosis 2 days from the start of the treatment **(Fig. 5E, Suppl. Fig 4E-F)**. However, co-administration of the pan-caspase inhibitor Z-VAD-FMK could efficiently suppress caspases activation **(Suppl. Fig. 4H)**, but did not rescue the cells from KHS101 cytotoxicity **(Fig. 5F, Suppl. Fig. 4G)**, indicating that cell death is not occurring solely as a result of apoptotic induction. Interestingly, no protection was observed also in presence of other cell death inhibitors of necroptosis or ferroptosis **(Suppl. Fig. 4I-J)**, suggesting that the metabolic alterations caused by KHS101 induce a cellular breakdown not rescuable by blocking the main cell death pathways. Furthermore, in order to investigate the impact of KHS101 *in vivo* we first tested the sensitivity of the tumor-forming mouse-derived NSCLC cell line LL2 **(Suppl. Fig. 5A-B)** and subsequently injected them in the tail vein of BL6 mice to evaluate lung metastasis growth upon KHS101 treatment. Lung tumors-bearing mice treated subcutaneously for 2 weeks with 6 mg/kg KHS101 twice a day did not show a significant difference in tumor area, as evaluated by IVIS measurement, but KHS101-treated mice showed prolonged disease-specific survival with increased survival time **(Suppl. Fig. 5C)**, in line with a non-significant trend for a reduced number of lung lesions, as macroscopically detected **(Suppl. Fig. 5D)**.

**Figure 5.**
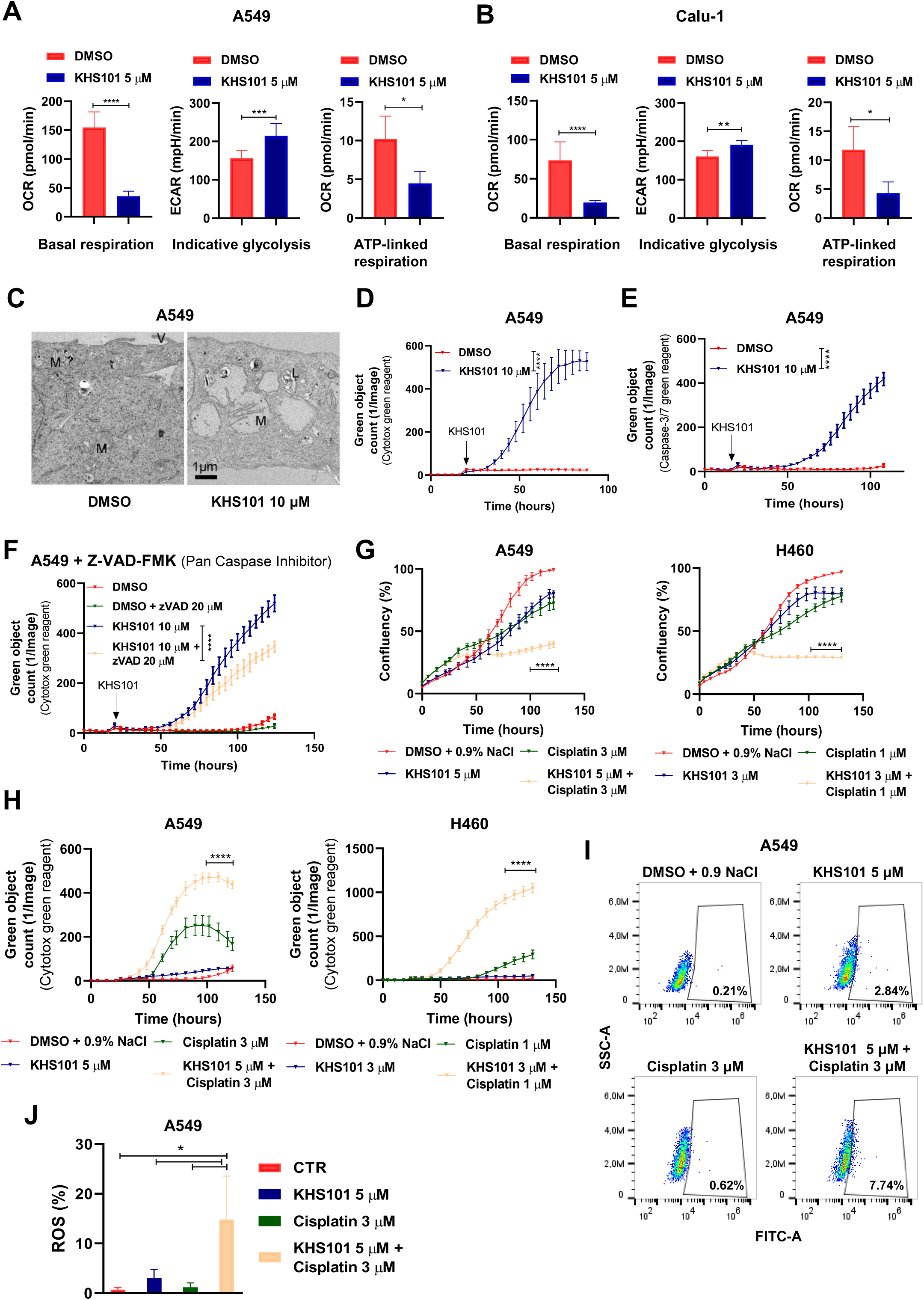
KHS101 induces a metabolic breakdown and death in NSCLC cells. Bar graphs showing quantification of basal respiration, indicative glycolysis and ATP-linked respiration in A549 (A) or Calu-1 (B) treated with KHS101 for 24 hours compared to control cells (DMSO). Bars (n=5) are average ± SD. P-values are from unpaired *t*-test. *<0.05, **<0.01, ***<0.001, ****<0.0001. C) Electron microscopy images of A549 treated 96 hours either with vehicle (DMSO) or KHS101 10 μM, (M: mitochondria, V: microvilli). D) Dead cell quantification as green object count using Cytotox green reagent (D) or caspse-3/7 reagent (E) of vehicle or KHS101 treated A549 cells. F) Dead cell quantification as green object count of A549 treated either with DMSO or KHS101 in combination with pan-caspase inhibitor Z-VAD-FMK. In D, E and F points (n=5) are average ± SD. P-value are from two-way ANOVA. ****<0.0001. G) Real-time proliferation curves of A549 and H460 treated with KHS101 and/or cisplatin compared to control cells. Death is shown as green object count in H). Points (n=5) are average ± SD. P-value are from two-way ANOVA. ****<0.0001. Percentage of ROS-positive cells in A549 cells treated with KHS101 and/or cisplatin for 24 hours are shown in FACS plots (I) and a graph bar (J). P-values are from one-way ANOVA. Scale bars (n=3) are average ± SD. *<0.05.

Finally, we investigated the *in vitro* effects of KHS101 in combination with cisplatin, one of the most commonly used chemotherapeutic drug used in relapsing NSCLC (29). Strikingly, a low cisplatin dose (1 or 3 µM) that alone was not able to strongly impair cell growth or survival, showed to induce a marked growth suppression **(Fig. 5G, Suppl. Fig. 6A)** and increase in cell death **(Fig. 5H, Suppl. Fig. 6B)** when combined with KHS101 (also at sub-lethal doses), resulting in a 2 to 4-fold reduced relative growth compared to cisplatin alone in three different cell lines **(Suppl. Fig. 6C)**. Cisplatin exerts its activity via upregulation of reactive oxygen species (ROS) (30) and by fluorescent ROS staining of treated cells we found that the combination of cisplatin and KHS101 synergistically increased ROS activation **(Fig. 5I-J)**. These data suggest that HSPD1 targeting can also induce cell death and increase the efficacy of chemotherapeutic treatments.

### Sensitivity to KHS101 is related to the metabolic state of the cells

We further profiled *in vitro* sensitivity to KHS101 by performing a screening on 26 different lung cancer cell lines, measuring cell viability 72 hours after treatment. Interestingly, none of the cell lines in the panel appeared to be completely resistant **(Fig. 6A, Suppl. Fig. 7A)**. KHS101-induced cytotoxicity was independent of the lung cancer type (NSCLC or SCLC) or NSCLC histological subtype **(Fig. 6B).** However, cells showed different grades of sensitivity towards the compound with IC_50_ values ranging from 2 to 14 μM **(Fig. 6C, Suppl. Fig. 7A)**. These results were independently validated comparing KHS101 sensitivity in the most sensitive and resistant cell lines based on percentage of inhibition of cell proliferation **(Suppl. Fig. 8A)**. In order to find possible determinants of drug resistance, cell lines with different genetic alterations were compared, but no particular genetic background was found as significantly protective **(Supp. Table 3)**. In order to identify possible molecular determinants of drug sensitivity, we performed a transcriptomic analysis on cell lines based on their IC_50_ values. HSPD1 mRNA expression correlated with IC_50_ values obtained from the screening **(Fig 6D)**. In order to have a stringent and genome-wide comparison, we used a 2-sided statistical approach. First, the 4 most resistant (NCI-H838, BEN, NCI-H1563 and SK-LU-1) and the 4 most sensitive NSCLC cell lines (H460, NCI-H1581, LOU-NH91 and A549) were grouped and the differentially expressed genes between the two groups were identified and ranked for fold difference **(Suppl. Table 4-5)**. Moreover, using available data from all the cells in the screening, we correlated the expression of each gene with the KHS101 IC_50_ values and ranked them based on Pearson metrics **(Suppl. Table 6-7)**. The results from these two analyses were overlapped **(Suppl. Fig. 8B)**, obtaining a list of potential up- and down-regulated genes in cells resistant to KHS101 **(Fig. 6E)**. The top up-regulated resistance-conferring gene was the transcription factor NLRC5, which acts as transactivator for the MHC class I genes (31) (some of which were also up-regulated in the analysis, like HLA-J or HLA-F). To validate this finding, we lentivirally induced NLRC5 overexpression in sensitive NSCLC cells **(Fig. 6F)**, and detected a significantly decreased cytotoxic effect of KHS101, indicating cellular resistance **(Fig. 6G, Suppl. Fig. 8C)**. Among the genes down-regulated in the resistant cell lines, the top hit was SLC6A8 **(Fig. 6E)**, a sodium chloride-dependent creatine transporter 1 (32). This evidence was validated by introducing a specific CRISPR/Cas9-based knock-out of SLC6A8, which increased cells viability of sensitive cell lines treated with KHS101 **(Fig. 6H)**. Since SLC6A8 is a creatine transporter we speculated that it could alter KHS101 sensitivity by changing the metabolic features of the cells. Analysis of a previously published metabolome profiling of NSCLC cell lines (33) revealed that cells that we determined to be the most KHS101 sensitive had a higher creatine level **(Fig. 6I)**, which was consistent with the results from the transcriptomic analysis (i.e. higher expression of the importer) suggesting a possible effect of creatine metabolism in the response to KHS101. Creatine can be phosphorylated by the cytosolic creatine kinase (C-CK) or by the mitochondrial isoenzymes (MtCK) to provide a rapid source of ATP (34) thereby fueling ATP-dependent cellular processes. To test whether differences in cellular creatine levels could be correlated with the cells’ metabolic state, we performed extracellular flux analysis of cells with higher (A549 and H460) and lower creatine (H838 and BEN) baseline levels (33) and found that cells with higher creatine had higher basal respiration **(Fig.6J)**. To further functionally prove this correlation, we inhibited creatine kinase activity with the specific inhibitor DNFB which decreased the basal respiration level of A549 cells **(Fig. 6K)**, concomitantly increasing the cells’ resistance to KHS101 as assessed by cell viability assays **(Fig. 6L)**. Taken together these findings suggest that a higher dependency on OXPHOS due to the alteration of creatine metabolism activity could determine the sensitivity to KHS101 of NSCLC cells.

**Figure 6.**
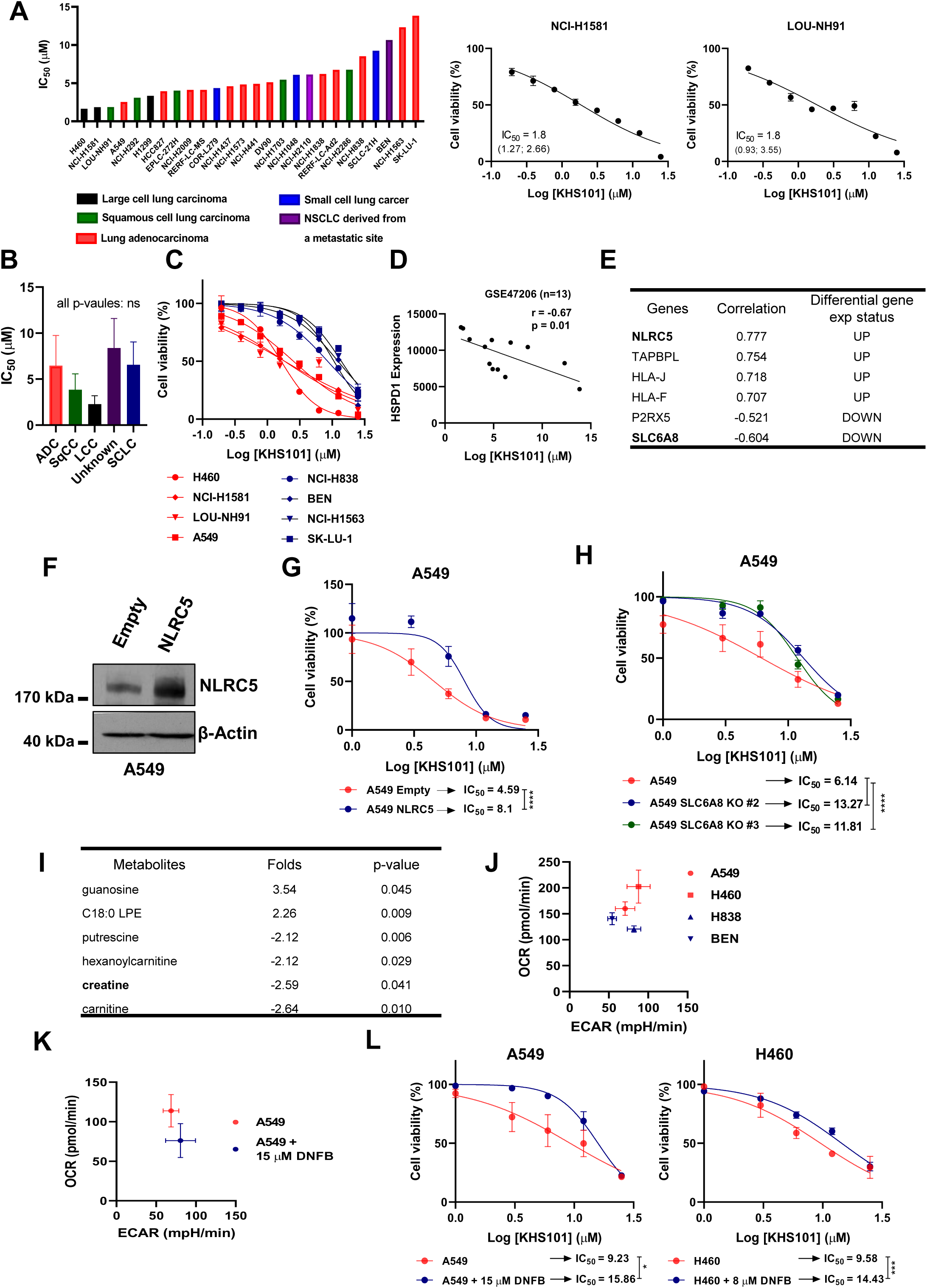
KHS101 sensitivity is related to NSCLC metabolic state. A) Bar graph showing IC_50_ values (μM) of all the 26 NSCLC cells belonging to CL-100 ProLiFiler screenin and dose-response curves (normalized to DMSO control) to KHS101 of 2 cell lines. Points (n=2) are average ± SD. IC_50_ values (μM) are shown with 95% confidence intervals. B) Bar graph showing the average IC_50_ for each NSCLC histotype and SCLC. Bars are average ± SD. P-values are from one-way ANOVA. C) Dose-response curves of the 4 more sensitive cell lines (H460, NCI-H1581, LOU-NH91, A549) compared to the 4 more resistant cell lines (NCI-H838, BEN, NCI-H1563, SK-LU-1) resulted from the screening. Points are average ± SD. D) Correlation between HSPD1 gene expression and KHS101 IC_50_ values in a panel of NSCLC cell lines from GSE47206 (n=13). E) Genes with most differential expression between KHS101-sensitive and KHS101-resistant groups (log_2_ fold change cut-off of 4 and p-value<0.05) F) Western blot analysis of NLRC5 protein level in A549 cells overexpressing NLRC5 or empty vector. β-Actin was used as loading control. G) Dose-response curves (normalized to DMSO control) of A549 expressing NLRC5 compared to empty control cells. Points (n=4) are average ± SD. IC_50_ values (μM) are shown. P-values are from two-way ANOVA. ****<0.0001. H) Dose-response curves (normalized to DMSO control) of A549 SLC6A8 KO compared to control cells. Points (n=4) are average ± SD. IC_50_ values (μM) are shown. P-values are from two-way ANOVA. ****<0.0001 I) Metabolite analysis between KHS101-sensitive and KHS101-resistant cells with a log_10_ fold change of 2 and p-value<0.05. J) Metabolic phenogram showing OCR (basal respiration) and ECAR (indicative glycolysis) values of creatine high (H460, A549) and creatine low (BEN, H838) cells. K) Metabolic phenogram showing OCR and ECAR levels in A549 treated for 24 hours with creatine-kinase inhibitor (DNFB). L) Dose-response curves (normalized to DMSO control) of A549 and H460 cells in presence of creatine-kinase inhibitor DNFB. Points (n=3) are average ± SD. IC_50_ values (μM) are shown. P-values are from two-way ANOVA. *<0.05, ***<0.001.

### KHS101 sensitivity depends on OXPHOS activity

To further investigate KHS101 resistance at a broader level, we performed a genome-wide CRISPR/Cas9-based dropout screening. A genetically barcoded whole-genome library (GeCKO v2) was transduced in A549 cells overexpressing the Cas9 nuclease. These cells were treated with two different KHS101 concentrations (10 and 15 μM) or DMSO as control **(Fig. 7A)**. None of the cells were able to survive the treatment with the 15 μM KHS101 dose, suggesting that is not possible to develop a complete resistance to this compound by the elimination of a single gene. By contrast, we could observe non-growing and morphologically altered cells surviving the 10 μM dose. After 2 weeks of survival (after all control pLKO-infected Cas9-A549 cells treated with the same KHS101 dose underwent complete cell death), the drug treatment was stopped to allow the selected cells to grow and be processed for the DNA preparation and the next-generation sequencing steps. The sequencing results on untreated cells indicated that it was possible to detect a total of 116700 sgRNAs (all gRNAs excluding those with no counts) present in the library, with a 97% representation, a level similar to previous screenings (25), while in the treated group this percentage dropped to 56%, as a result of the drug selection. Hits were ranked and gene ontology analysis on the 20 top hits (**Supplementary Table 7**) indicated that the genes whose knockout increased cellular resistance to KHS101 were mainly involved in mitochondria structure and activity **(Fig. 7B)**. The COX5B gene, a subunit of the cytochrome c oxidase, was the ‘top hit’. In order to independently validate this finding, we knocked out COX5B by two independent gRNAs in A549 cells, and in both cases we found the emergence of a more KHS101-resistant phenotype **(Fig. 7C)**. COX5B knock-out cells presented a reduction in their basal respiration and ATP-linked respiration **(Fig. 7D)**, strengthening the notion that KHS101 sensitivity is linked with cellular oxidative activity, in line with the results obtained from creatine metabolism. Conversely, blocking glycolysis by 2-deoxy-D-glucose (2-DG) treatment significantly enhanced KHS101 sensitivity in different cell lines **(Fig. 7E),** further confirming that cancer cells respond to KHS101 proportionally to their OXPHOS-dependence. Therefore, HSPD1 targeting induces a mitochondrial metabolic breakdown in NSCLC, which results in loss of growth potential and cell death, and the small molecule KHS101 is particularly active on OXPHOS-dependent cells.

**Figure 7.**
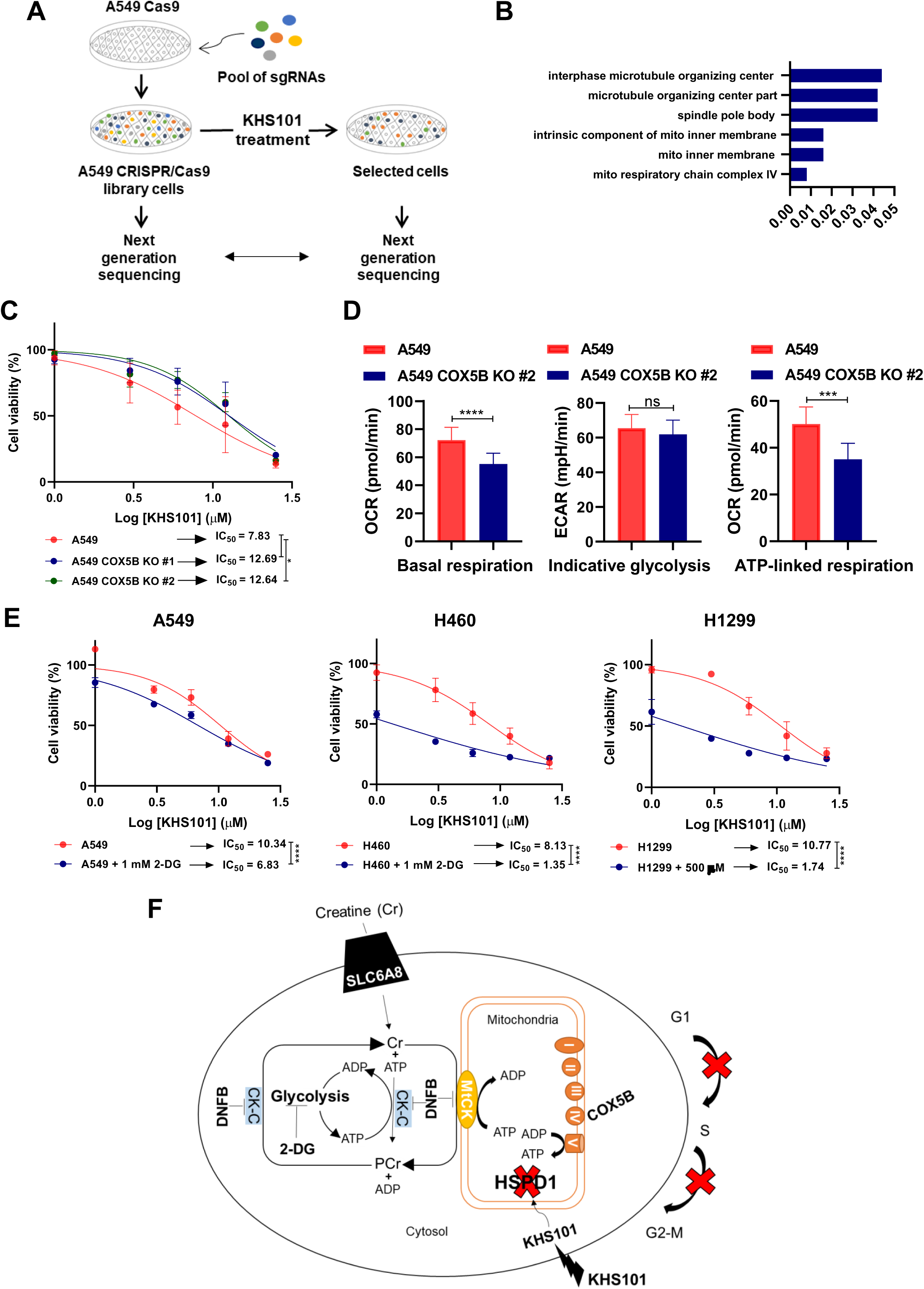
Mitochondrial metabolism influences NSCLC cells response to KHS101. A) Scheme showing the genome-wide CRISPR-Cas9 screening platform employed to identify KHS101 resistance genes. B) Gene ontology (GO) analysis of the top 20 candidates identified from the knockout screen in cells after treatment with KHS101. GO was performed with the cellular component 2017b gene set library in Enrichr tool. C) Dose-response curves to KHS101 (normalized to DMSO control) of A549 cells overexpressing two gRNAs to knockout COX5B, compared to parental A549. Points (n=4) are average ± SD. IC_50_ values (μM) are shown. P-values are from two-way ANOVA. *<0.05. D) Bar graphs showing quantification of basal respiration, indicative glycolysis and ATP-linked respiration of A549 COX5B knockout cells compared to parental cells. Bars (n=5) are average ± SD. P-values are from unpaired *t*-test. ***<0.001, ****<0.0001. E) Dose-response curves (normalized to DMSO control) of A549, H460 and H1299 cells in presence of glycolysis inhibitor 2-deoxy-D-glucose (2-DG). Points (n=3) are average ± SD. IC_50_ values (μM) are shown. P-values are from two-way ANOVA. ****<0.0001. F) Schematic representation of the metabolic alteration that occur in NSCLC cells upon HSPD1 loss.

## DISCUSSION

The identification of novel targets and the development of more effective drugs is of paramount importance for improving the clinical management of NSCLC. Metabolism has recently gained much attention in the oncological field and anti-metabolism strategies have been proposed to reduce the lethal effects of NSCLC (35-37). We report here heat shock protein HSPD1 as a theranostic metabolic marker for NSCLC, whose elimination can have profound effect on outcome of the patients.

Heat shock proteins (HSPs) are molecular chaperones categorized according to their molecular sizes into small HSPs and large HSPs (16). HSPs can be highly expressed in cancer cells, where they promote cell proliferation, metastasization and drug resistance, HSPD1 is an ATP-dependent HSP mainly localized in the mitochondria, playing an essential role in guaranteeing the correct folding of the mitochondrial-imported proteins (38) and some data are available on its role as a stemness/metastasis regulator in other cancers (22,39,40). In lung cancer, *HSPD1* has been previously found highly expressed in NSCLC tissues (41) and identified as a predictive marker for survival in both smokers and non-smokers patients (42). Our findings confirm that HSPD1 is ubiquitously expressed at a high level (as detectable by IHC in all samples) and is associated with adverse prognosis, adding the notion that its expression is essential for NSCLC survival (fitness gene), making it a very attractive target for future therapy. In order to more functionally explore this possibility we evaluated the effect of HSPD1 targeting achieved by either shRNA-mediated knockdown or by a specific small molecule. In shRNA experiments, the level of growth suppression measured for each shRNA sequence was generally proportional to the degree of HSPD1 knockdown. Importantly, even if a complete protein elimination was not achieved, the cell growth blockage obtained by the best performing shRNA (#48) was durable (up to one month) and very effective both *in vitro* and *in vivo* in all the cell lines investigated. When cellular metabolic changes were measured, cells with HSPD1 knockdown presented a strong reduction in basal respiration and impaired capacity in synthesizing ATP though OXPHOS, in line with the role of HSPD1 in the (re)folding of mitochondrial-imported oxidative respiration proteins (43). The activity of HSPD1 is therefore essential for maintaining NSCLC metabolic fitness and its loss causes a profound energetic breakdown affecting the ability of the cancer cells to divide and expand.

An emerging way of chemically disrupting the functionality of HSPD1 is the treatment with the synthetic small compound KHS101. This compound was originally identified through a phenotypical screen as an inducer of neuronal differentiation both in vitro and *in vivo* (44). KHS101 promotes HSPD1 aggregation thereby altering the metabolic activity of glioblastoma cells (23). In NSCLC, we observed that KHS101-treated cells were very severely growth arrested, in analogy to what was observed with the HSPD1 knockdown approach. OXPHOS activity was also efficiently repressed, even with low micromolar KHS101 concentrations, starting from a few hours after treatment. In contrast to HSPD1 knockdown, KHS101-treated cells increased glycolysis, suggesting that they might try to compensate for the loss of ATP production. This increase is, however, not sufficient for the cells to gain resistance and survive the treatment. Indeed, KHS101-treated cells underwent significant cell death, probably due to the formation of irreversible HSPD1 multi-aggregates coagulated by KHS101, as shown (23). Notably, co-administration of specific inhibitors of death pathways did not rescue KHS101 treated cells, indicating that the occurring death is strong, multifactorial, and irreversible, all features of crucial importance for a novel anti-cancer therapeutic strategy.

In order to better understand the biology of HSPD1 targeting, two independent unbiased screenings were carried out: a differential gene expression analysis on KHS101 sensitive and resistant cells and a genome-wide CRISPR/Cas9-based dropout screening on drug treated cells. The first approach identified a biomarker of increased sensitivity, the creatine importer SLC6A8, which lead us to discover OXPHOS dependency as condition for better KHS101 efficacy. Moreover, a potential role in determining KHS101 resistance of the MHC class I genes transactivator NLRC5 as been here identified. Based on its importance on inflammatory processes in tumors (31), the molecular details of this link should be in the future determined. The second approach came independently to a similar and converging conclusion, by identifying mitochondrial metabolism genes, and in particular the subunit of the cytochrome c oxidase complex (COX5B), as enriched in cells surviving KHS101. All these evidences highlight the targeting of mitochondrial metabolism as an effective therapeutic strategy to achieve a complete, irreversible elimination of NSCLC.

The occurrence of drug-overcoming mutations is a typical feature of NSCLC, which has unfortunately so far limited the success of targeted therapies (45). Here, noteworthy, not a single cell from the whole-genome CRISPR/Cas9 library was able to survive the KHS101 selection when administered at the highest dose (15 μM). At the lowest dose (10 μM), a small population of cells was able to persist, but could not grow (and be processed for sequencing) until the drug was entirely removed from the experiment. This is a very important observation in line with the observed inability of death pathways inhibitors to protect *in vitro* from KHS101, and in line with the data from the two independent fitness analyses on *HSPD1* gene and with the drug screening from large collections of NSCLC cell lines of various genetic backgrounds, which confirmed that no single genetic alteration can guarantee a complete protection from the treatment with KHS101. The sensitivity towards KHS101 is also independent from the lung cancer type or NSCLC histotype, suggesting that a HSPD1 targeting approach might be valuable also for other cancer types. Another aspect of fundamental importance to consider for NSCLC drug development is the potential successful combination with chemotherapy, as chemotherapy still represents the backbone of therapeutic regimens, especially in late-stage lung cancer patients (46-48). Even if the data here presented will require future dedicated experimentation, the results are encouraging, as KHS101 showed to act synergistically with cisplatin *in vitro*, enhancing the production of reactive oxygen species and reducing the dose of cisplatin needed to slow down or kill the cancer cells. All these evidences reinforce the potential relevance of designing and developing HSPD1-targeting drugs for NSCLC therapy. This goal, however, is not likely to be achieved by KHS101 itself, which this study found active at relatively higher concentrations. KHS101 therefore stands out as a rather excellent tool compound for accelerating discoveries for HSPD1 targeting, but novel drugs (or modifications of the same compound) with improved affinity will need to be designed and tested, for instance by taking advantage of emerging targeting techniques based on induction of selective intracellular proteolysis (49,50). The activity of these drugs could be in the future tested beyond NSCLC, as HSPD1 appears to be a fundamental oncoprotein also in other contexts (39,40).

In conclusion, we showed that HSPD1 elimination interferes with NSCLC metabolic activity causing a strong OXPHOS-dependent energetic breakdown, which the cancer cells fail to overcome, highlighting HSPD1 as a powerful theranostic marker for improving lung cancer therapy.

## AUTHORS’ CONTRIBUTIONS

Conception and design: B.P, P. C.; Development of methodology: B.P, V.R, P.G, A.S, L.P, A.S, S.M, S.Z, C.P, S.U, D.M, H.S, H.W, P. C.; Acquisition of data (provided animals, acquired and managed patients, provided facilities, etc.): F.N, M.V, S.U, D.M, M.S; Analysis and interpretation of data (e.g., statistical analysis, biostatistics, computational analysis):B.P, V.R, P.C; Writing, review, and/or revision of the manuscript: B.P, H.W, P.C; Administrative, technical, or material support (i.e., reporting or organizing data, constructing databases): V.R, P.G, A.S, S.M, S.Z, P.C, S.U, D.M, M.S, H.W, P.C. Study supervision: P. C. The author(s) read and approved the final manuscript.

---

**Suppl. Figure 1.**
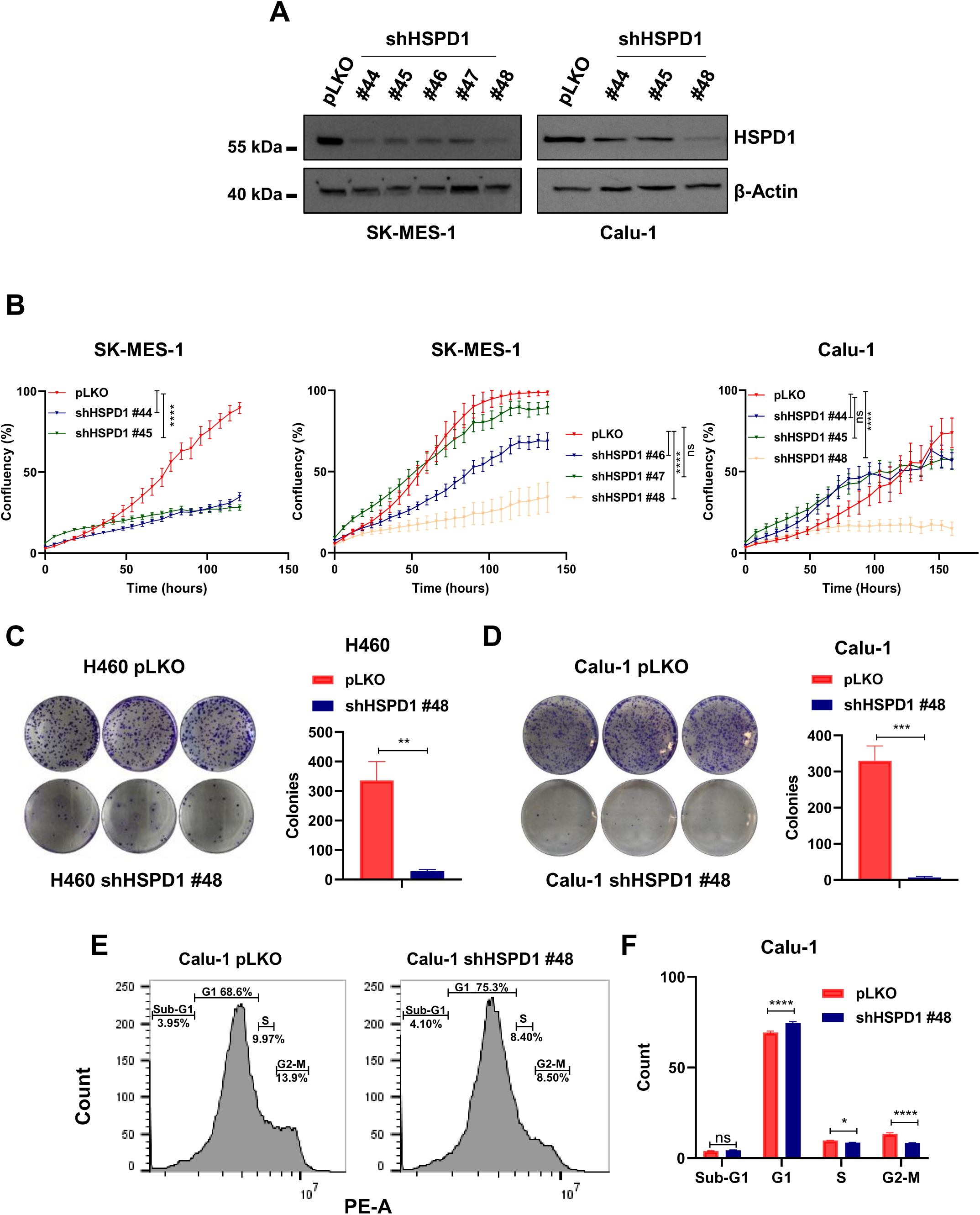
A) Western blot analysis of HSPD1 protein level in SK-MES-1 and Calu-1 cells upon infection with respectively 5 or 3 independent shRNAs (#44, #45 #46, #47 and #48) targeting HSPD1 compared to scramble-infected cells (pLKO). β-Actin was used as loading control. B) Real-time proliferation curves of SK-MES-1 and Calu-1 infected with non-targeting pLKO or shHSPD1. Plotted is cells’ confluency over time. P-values are from two-way ANOVA. Points (n=5) are average ± SD. ****<0.0001. Colony formation of H460 (C) and Calu-1 (D) cells infected with pLKO or shHSPD1, stained with crystal-violet and quantified in triplicates. Scale bars are average ± SD. P-values are from unpaired *t*-test. **<0.01, ***<0.001. E) FACS plots of Calu-1 cells infected with pLKO or shHSPD1 and stained with PI for cell cycle analysis. F) Bar graph showing the % of cells in each cell cycle phase of Calu-1 cells upon infection with pLKO or shHSPD1. P-values are from two-way ANOVA. Scale bars (n=3) are average ± SD. *<0.05, ****<0.0001.

**Suppl. Figure 2.**
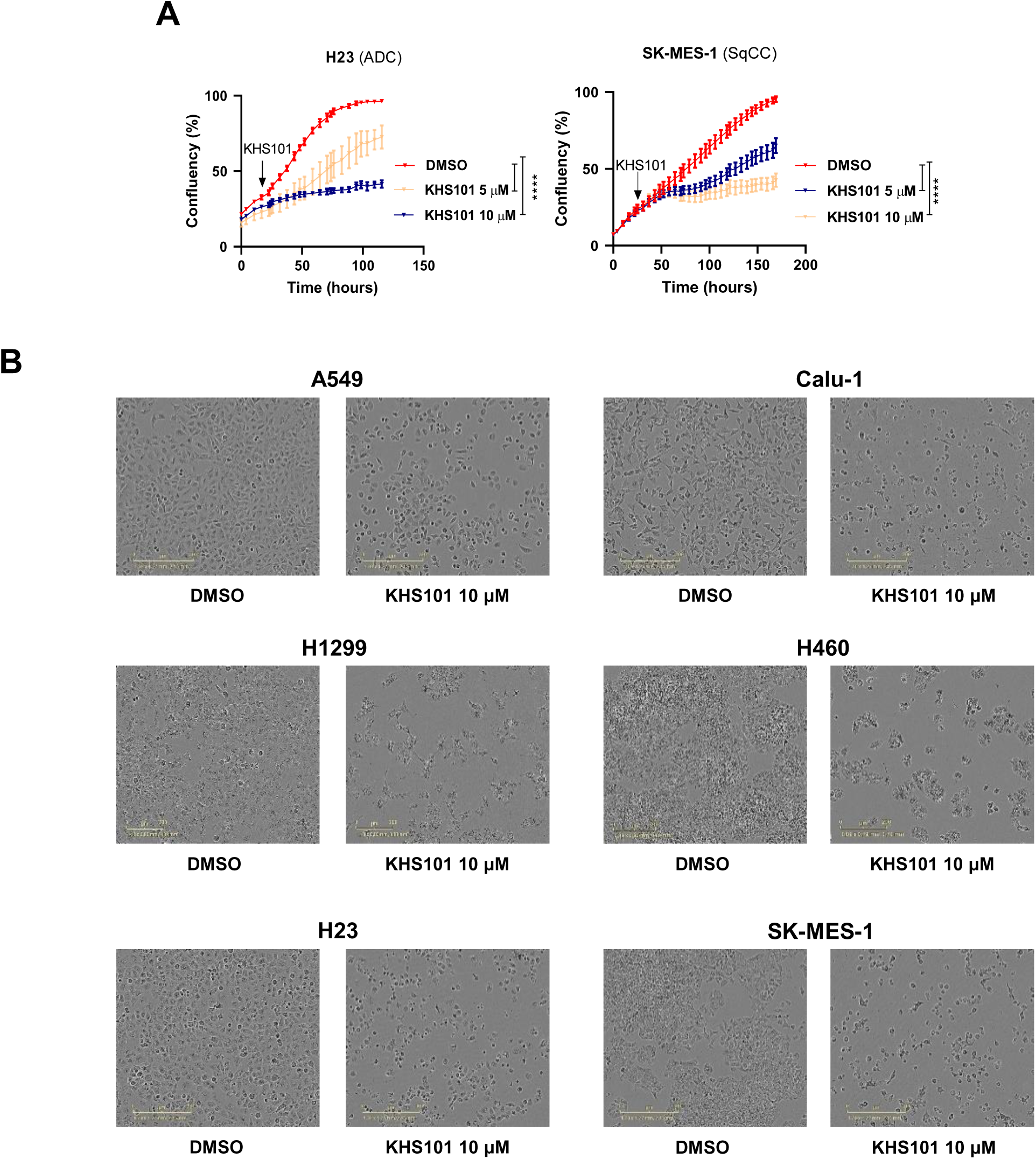
A) Real-time proliferation curves of H23 and SK-MES-1 treated either with vehicle (DMSO) or KHS101. P-values are from two-way ANOVA. Points (n=5) are average ± SD. ****<0.0001. B) Images of cells (A549, Calu-1, H1299, H460, H23 and SK-MES-1) treated with DMSO or KHS101 10 μM (after 5 days of treatment).

**Suppl. Figure 3.**
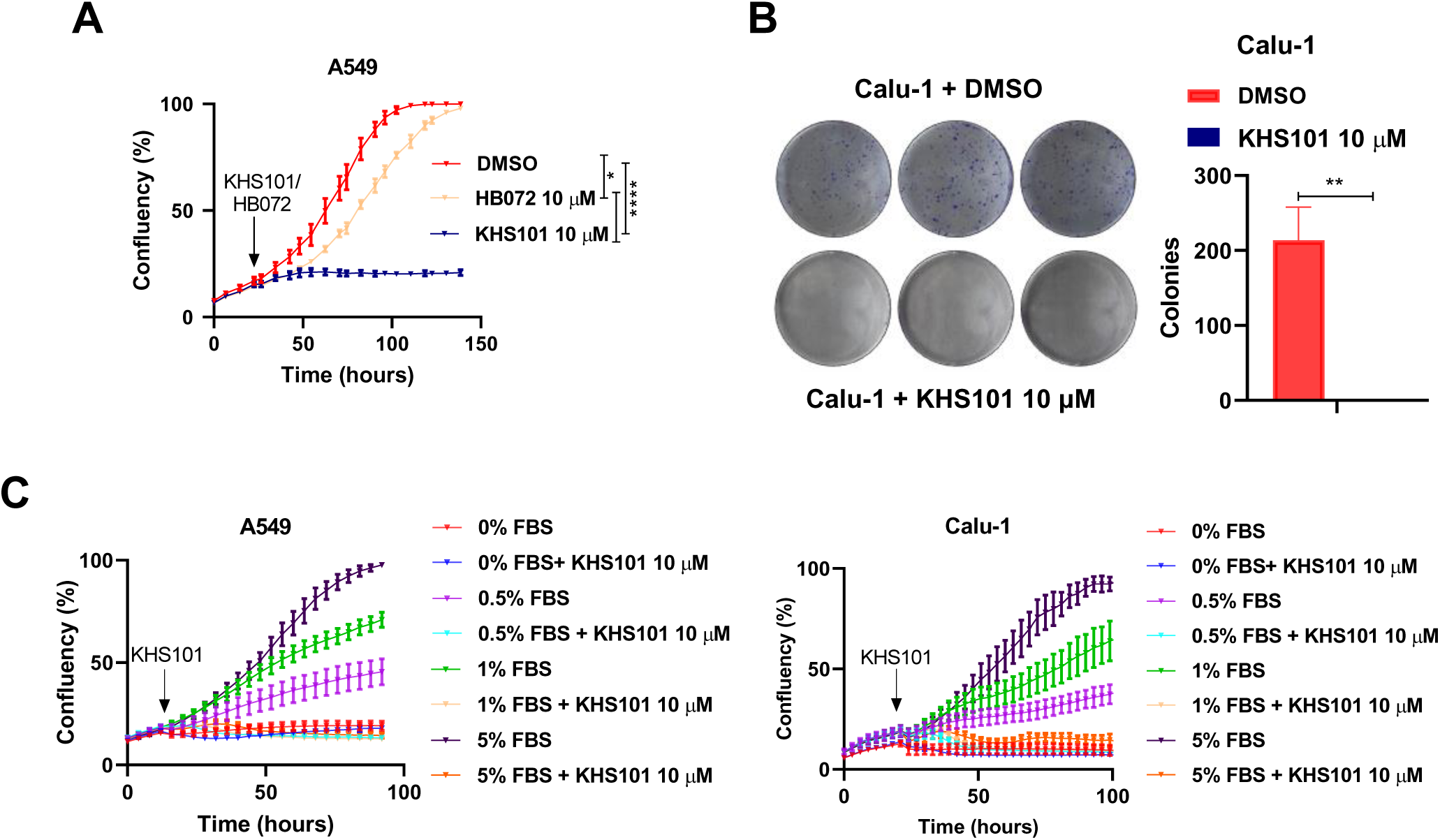
A) Real-time proliferation curves of A549 cells treated either with KHS101 or with the corresponding inactive KHS101 analog (HB072) compared to vehicle-treated cells. Points (n=5) are average ± SD. P-values are from two-way ANOVA. *<0.05, ****<0.0001. B) Colony formation of Calu-1 treated for 5 days with KHS101 or vehicle and then left to grow in drug-free media, stained with crystal-violet and quantified in triplicates. Scale bars are average ± SD. P-values are from unpaired *t*-test. **<0.01. C) Real-time proliferation curves of A549 and Calu-1 treated with KHS101 in presence of different concentration of FBS in the media. Points (n=5) are average ± SD.

**Suppl. Figure 4.**
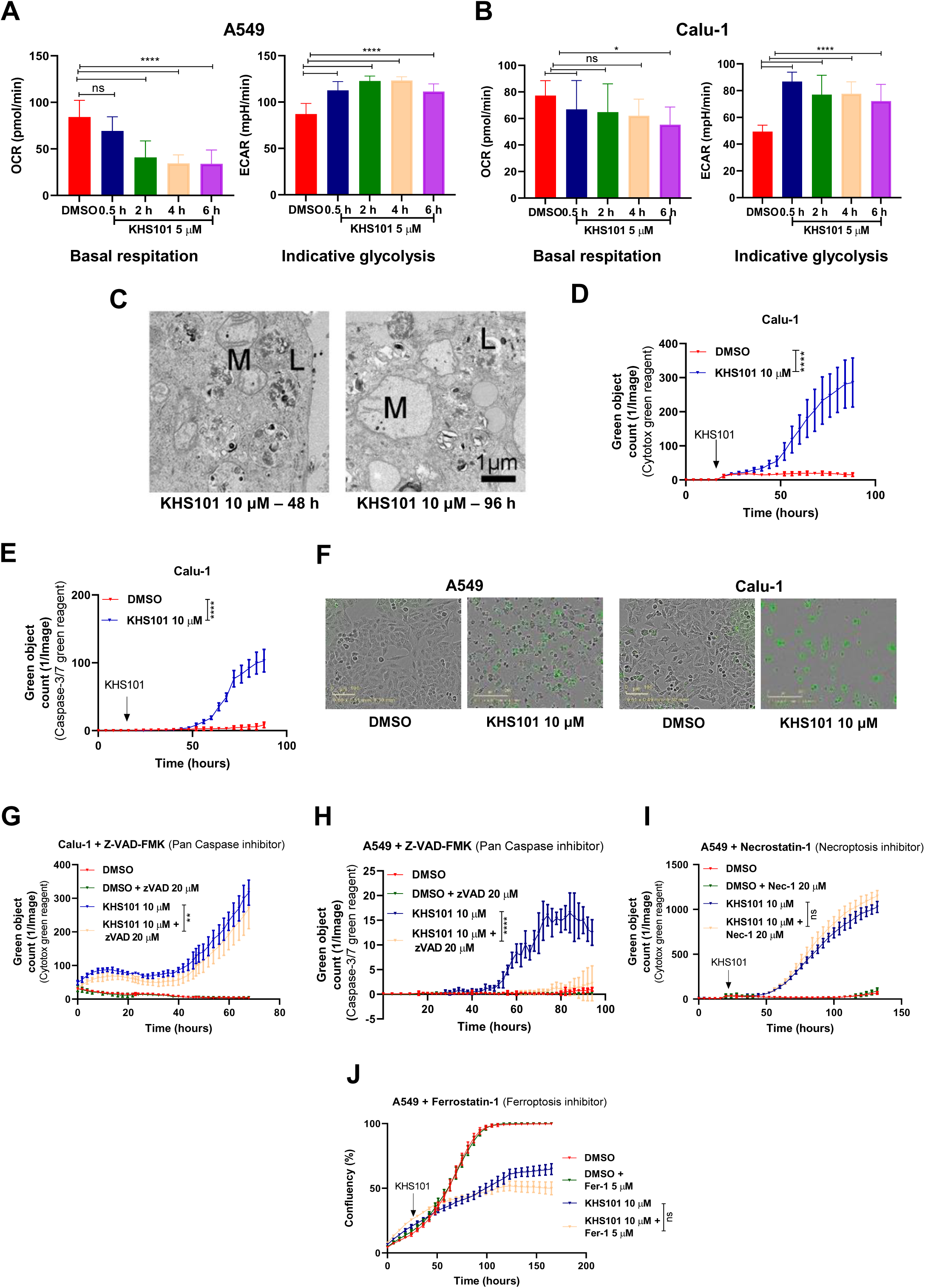
Quantification of OCR (basal respiration) and ECAR (indicative glycolysis) of A549 (A) and Calu-1 (B) cells treated for 0.5 h, 2 h, 4 h and 6 h compared to control cells. Bars (n=5) are average ± SD. P-values are from unpaired *t*-test. *<0.05, ****<0.0001. C) Electron microscopy images of A549 treated either for 48 or 96 hours with KHS101 10 μM. L-lysosomes, M-mitochondria. Dead cell quantification as green object count using Cytotox green reagent (D) or caspse-3/7 reagent (E) of vehicle or KHS101 treated Calu-1 cells. F) Images of A549 and Calu-1 cells treated with vehicle or KHS101 and stained with caspase-3/7 reagent. G) Dead cell quantification as green object count of Calu-1 treated either with DMSO or KHS101 in combination with pan-caspase inhibitor Z-VAD-FMK. H) Dead cell quantification as green object count using caspase3/7 green reagent of A549 treated either with DMSO or KHS101 in combination with pan caspase inhibitor Z-VAD-FMK. I) Dead cell quantification as green object count of A549 treated either with DMSO or KHS101 in combination with necroptosis inhibitor Necrostatin-1. J) Real-time proliferation curves of A549 treated either with DMSO or KHS101 in presence of ferroptosis inhibitor Ferrostatin-1. In D, E, G, H, I and J points (n=5) are average ± SD. P-values are from two-way ANOVA. **<0.01, ****<0.0001.

**Suppl. Figure 5.**
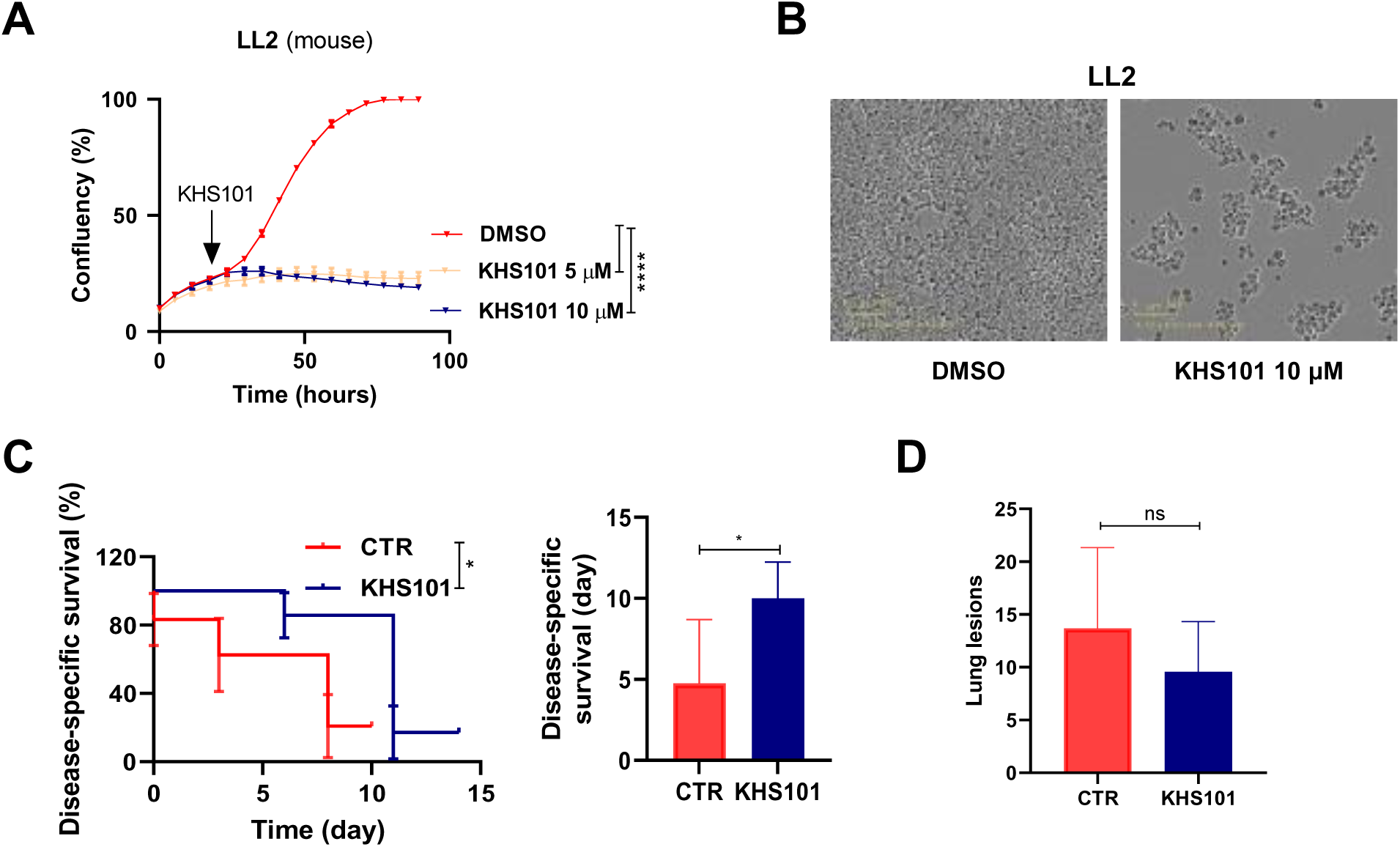
A) Real-time proliferation curves of mouse cell lines LL2 treated either with vehicle (DMSO) or KHS101. Points (n=5) are average ± SD. P-values are from two-way ANOVA. ****<0.0001. B) Images of Ladi3.1 and LL2 treated with DMSO or KHS101 10 μM. C) Kaplan–Meier disease-specific survival analysis of C57BL/6 mice treated with 6 mg/kg KHS101 or vehicle (5% (v/v) Ethanol - 15% (w/v) Captisol solution) for 2 weeks. P-value is Log-rank test. *<0.05. Disease-specific survival time (shown on the right panel) is calculated starting from the first appearance of bioluminescence. Bars are average ± SD. P-value is unpaired *t*-test. *<0.05.

**Suppl. Figure 6.**
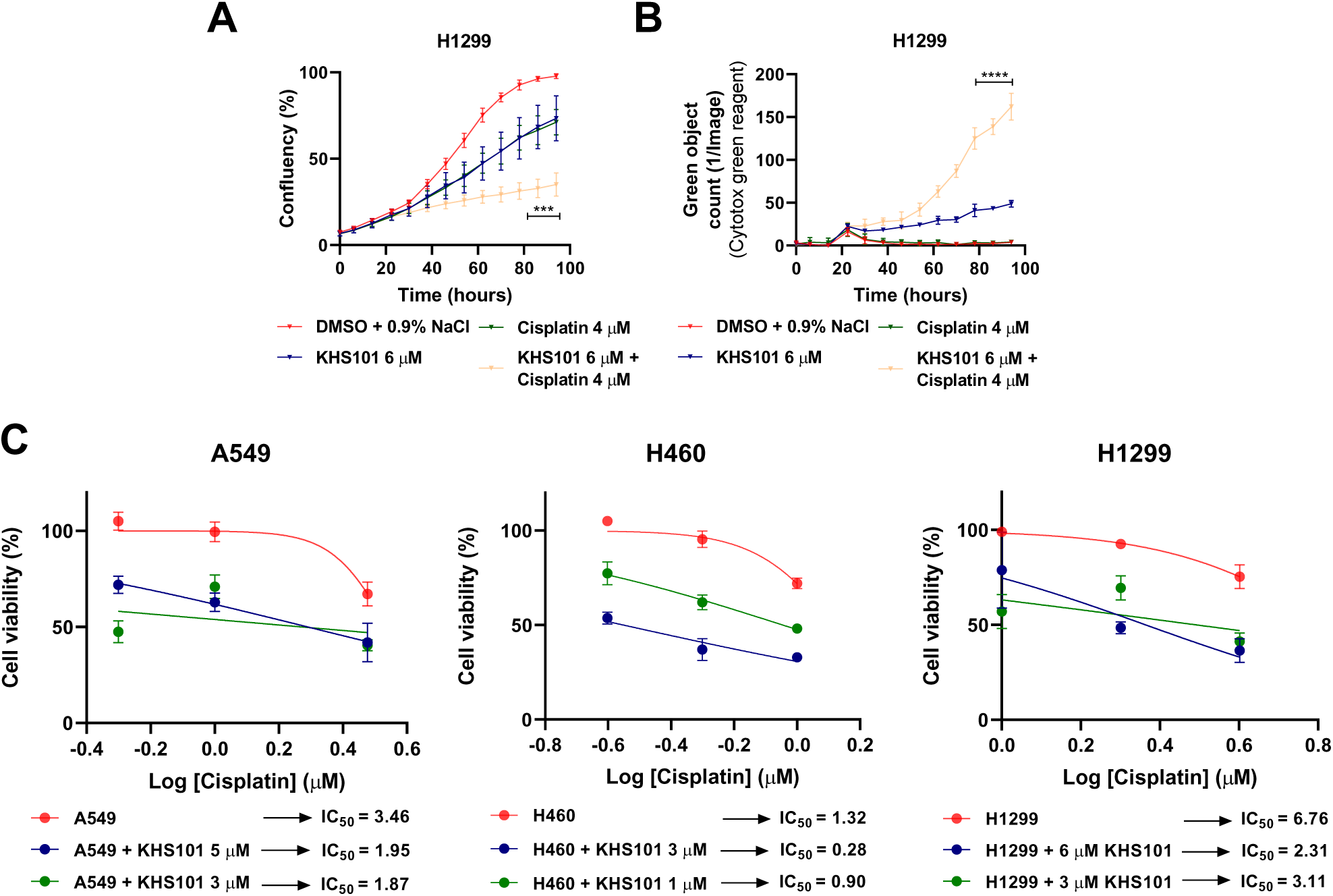
Real-time proliferation curves (A) and dead cell quantification (B) of H1299 cells treated with KHS101 and/or cisplatin at indicated doses compared to control cells. Points (n=5) are average ± SD. P-value are from two-way ANOVA. ***<0.001, ****<0.0001.C) Dose-response curves to cisplatin of A549, H460 and H1299 cells treated with different concentration of KHS101. IC_50_ values (μM) are shown. Points (n=5) are average ± SD.

**Suppl. Figure 7.**
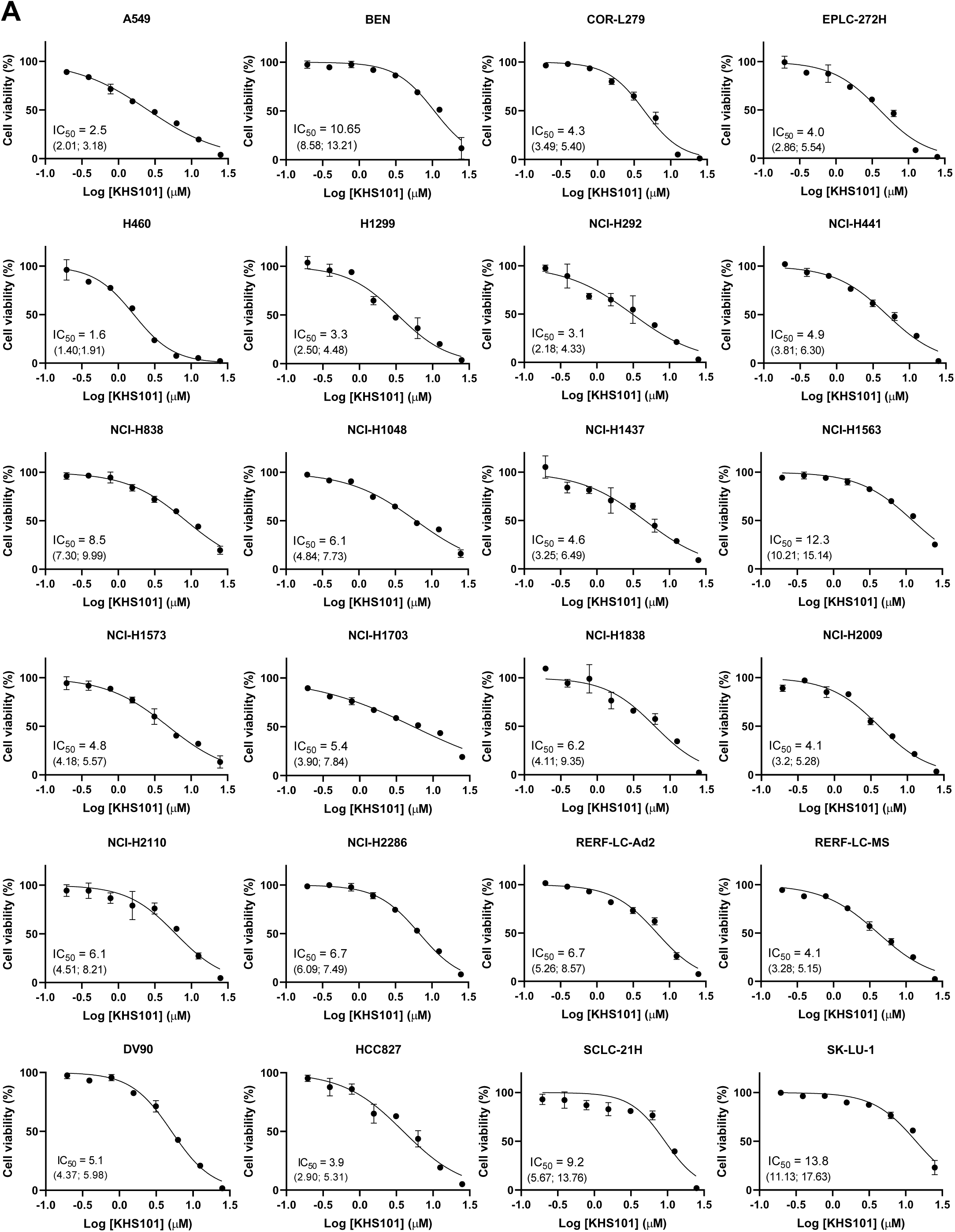
A) Dose-response curves (normalized to DMSO control) to KHS101 of the remaining 24 NSCLC cell lines belonging to CL-100 ProLiFiler screening. Points are average ± SD. Their IC_50_ values (μM) are shown with 95% confidence intervals.

**Suppl. Figure 8.**
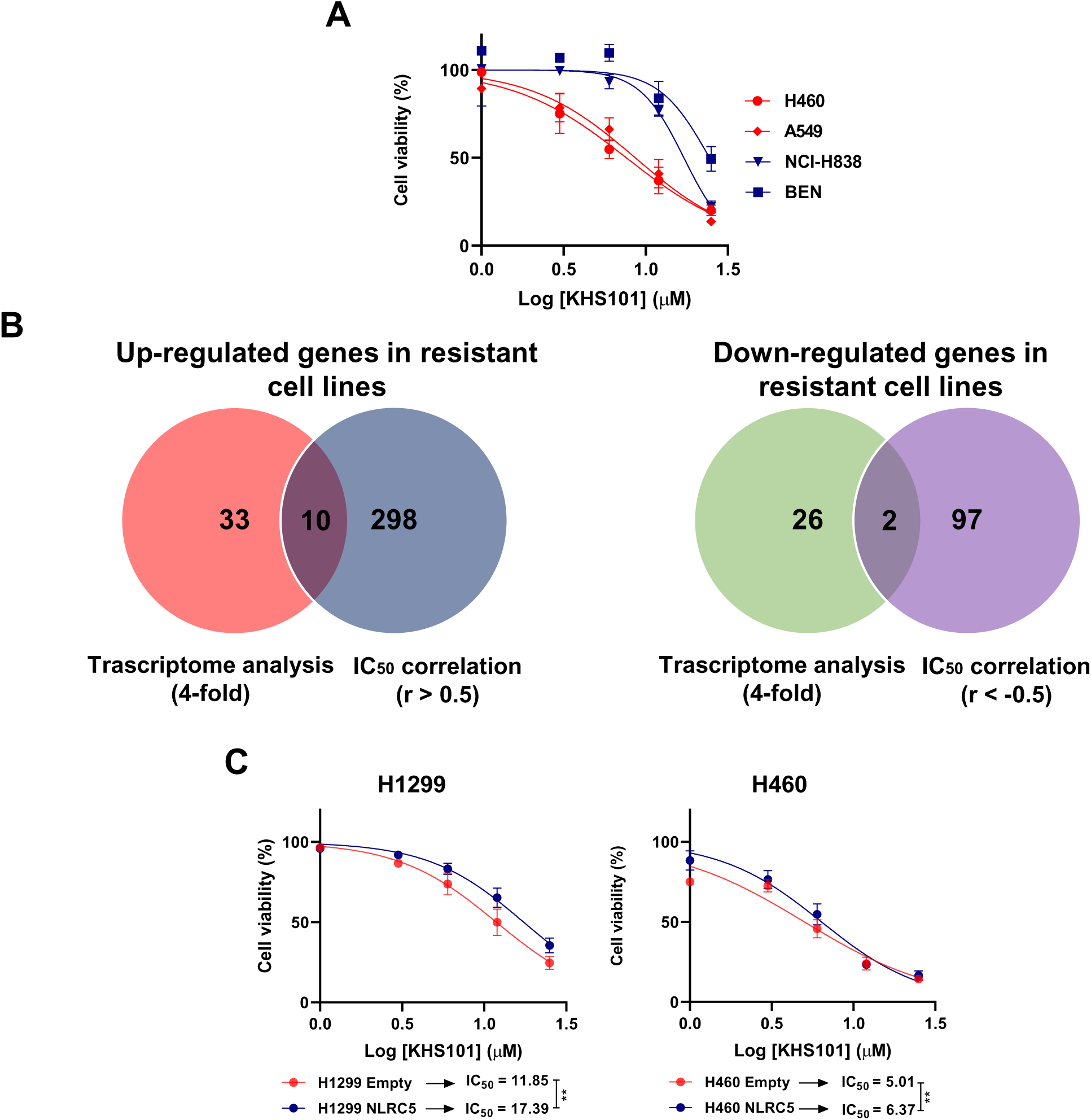
A) Dose-response curves of 4 cell lines (H460, A549, NCI-H838 and BEN) as validation of the CL-100 ProLiFiler screening. Points (n=5) are average ± SD B) Venn diagrams showing overlap of up- or down-regulated genes in resistant cells identified in the transcriptome analysis and IC_50_ correlation analysis. C) Dose-response curves (normalized to DMSO control) of H1299 and H460 overexpressing NLRC5 compared to empty control cells. Points (n=4) are average ± SD. IC_50_ values (μM) are shown. P-value is from two-way ANOVA. **<0.01.

## SUPPLEMENTARY METHODS

### Lentiviral transduction

Plasmids for HSPD1 knock down (TRCN0000029444, TRCN0000029445, TRCN0000029446, TRCN0000029447 and TRCN0000029448 for human cell lines) are from Sigma. Scrambled pLKO.1 (referred to as pLKO) was used as non-targeting control. NLRC5 expression vector (EX-A3335-Lv105) and control vector (Ex-Neg-LV105b) are from GeneCopoeia. Plasmids containing RNAs for COX5B (gRNA sequence #1 is ACTTCGCGGAGCTGGAACGC; gRNA sequence #2 is CAGCCAAAACCAGACGACGC) and SLC6A8 (gRNA sequence #2 is ACGGGGCCGTCGCCCTTGGC; gRNA sequence #3 is GCCCTTACCATGCAGACCAG) knock-out (pLENTI-CRISPR-V2) are from Genscript. For production of lentiviral particles, 293T cells were transfected with 8 μg knock-down/expression vectors and 2 μg of packaging vectors (pMDL, pVsVg and pRevRes) in complex with 24 μg PEI (Polysciences) in 0.9% NaCl. After 48 hours, supernatant was collected, centrifuged and filtered. For transduction, 150,000 cells were seeded in a 6-well plate and infected in presence of 8 μg/mL polybrene (Sigma). Selection was done with 3 μg/mL puromycin (Sigma) and cells were maintained in 1 μg/mL puromycin.

### Western blot analysis

For proteins isolation, cells were lysed in RIPA buffer and quantified using Pierce BCA kit (Thermo-Fisher). For cytosolic/mitochondrial fractionation Cytochrome c Release Assay Kit (abcam, ab65311) was used. Proteins lysates (10–20 μg) were resolved on 5%-12% SDS–PAGE gels and transferred to PVDF membrane (Thermo-Fisher). Membranes were blocked in 5% Milk (BioRad) or 5% BSA (Sigma) in 1X TBST and incubated overnight at 4°C in primary antibodies. Membranes were then washed with 1X TBST and incubated with secondary antibodies (Southern Biotech) for 1 hour. The membranes were developed with ECL reagent (Thermo Fisher) on to X-ray films (Thermo-Fisher) using the chemiluminescence imager, AGFA CP100. Rabbit anti-HSPD1 (ab46798, 1:20000) and rabbit anti-NLRC5 (ab117624, 1:1000) antibodies were purchased from Abcam; mouse anti-TOMM20 (H00009804-M01, 1:1000) was purchased from Abnova; anti-β-Actin (8H10D10) HRP conjugate (1:10000) was purchased from Cell Signaling

### Colony formation assay

To assess clonogenic ability of knockdown cells or KHS101 treated cells, a colony formation assay was performed. Control (pLKO) and knockdown cells were plated in triplicates at a low density (1,500 cells/well) in a 6-well plate in triplicates and they were allowed to grow until they formed visible colonies. To assess, instead, the clonogenic ability upon KHS101 treatment, 200,000 cells were seeded in triplicates in 6-well plates and then treated with 10 μM KHS101 for 5 days. For KHS101 treated cells the media was replaced with normal media, whereas the DMSO cells where collected, counted and seeded in triplicates at low density (1,500 cells/well) and they were allowed to grow until they formed visible colonies. Cells were washed with 1X PBS and fixed in 10% formalin (Sigma) for 5 minutes. After washing, colonies were stained with 0.05% crystal violet solution (Biomatik, CAS 548629) for 30 minutes and washed twice with deionized water. Colonies were photographed and counted.

### FACS analysis

Cell cycle analysis was performed using the Propidium iodide (PI; Sigma) staining. 50,000 cells were seeded in triplicates in 12-well plates and incubated overnight. Samples were prepared as described in (51). For ROS staining, 50,000 cells were seeded in 12-well plates and the day after they were treated with KHS101 in combination with cisplatin. After 24 hours, samples were washed and stained for 30 minutes at 37°C with CM-H2DCFDA 5 μM (Thermo Fisher, C6827). Then they were collected in FACS tubes, washed and re-suspended in FACS buffer (2% FBS, 5 mM EDTA in 1X PBS). Samples were run on Cytoflex FACS machine (Beckman) and data were analyzed using FlowJo software v10.6.

### Immunohistochemistry (IHC)

HSP60 (HSPD1) immunohistochemistry was performed on 20 consecutive NSCLC lung surgical specimens using an automated platform (BenchMark, Ventana Medical Systems, Roche, Basel, Switzerland). Briefly, samples were pretreated for 36 minutes with antigen retrieval ULTRA CC1 then they were incubated for 32 minutes at 36°C with HSP60 (HSPD1) (AbTA800758, Clone OTI3A2, 1:100 dilution, ORIGENE, Rockville, US) primary antibody. The use of retrospective LC tissues for immunohistochemical study was approved by the Research Ethics Committee of the San Luigi Hospital/University of Turin (approval n.26 dated January 18^th^ 2019). Informed consent was obtained from all patients. Samples have been anonymized by a staff member not involved in the study and all patient information are made unavailable to the investigators.

### Generation of CRISPR/Cas9 lentiviral library and CRISPR screen

CRISPR/Cas9 lentival library was generated as described in (25). A549 Cas9 cells were transduced with serial dilutions of a virus to find the MOI of ∼0.3. A549 Cas9 cells were transduced with lentiviral Human GeCKO v2 knockout pooled library part A and part B (a gift from Feng Zhang (52), Addgene # 1000000049) at MOI of 0.3 in the presence of 10 μg/mL of polybrene for 24 hours, then replaced the virus medium with fresh growth medium and continued to culture the cells for 48 hours. The cells were selected with 50 μg/mL of puromycin for 3 days. Then 4 million cells were seeded in 10-cm dishes and treated the day after either with vehicle or with 15/10 μM KHS101. A549 Cas9 pLKO were used as positive control. After 2 weeks, the media containing the drug was replaced with normal media to allow the cells to grow again. 40 million cells were then collected for genomic DNA isolation. Next generation sequencing was performed on the Illumina HiSeq 2500 platform in Deep Sequencing Facility of TU Dresden. The raw FASTQ files were analyzed with MAGeCK-VISPR (53).

### Genomic DNA isolation and PCR amplification

Genomic DNA was extracted with NucleoSpin® Blood XL (Machery Nagel # 740950.50) according to the manufacturer’s protocol. The first round PCR of Next Generation Sequence (NGS) is performed with 26 separate 100 μL redundant reactions, each containing 5 μg of DNA, 50 μL Q5® Hot Start High-Fidelity 2X Master Mix (NEB # M0494L), and 3 μL of a 10 μM solution of each primer (P5 and P7). The PCR was performed as described in (25).

### Transmission electron microscopy

A549 cells were treated either with vehicle (DMSO) or KHS101 10 μM for 48 and 96 hours. After treatment the cells were fixed with 2% glutaraldehyde in 0.04 M phosphate buffer pH 7.4 for 60 minutes, scraped off, collected in a tube and post-fixed with the same fixative for 24 hours. The cells were then washed in PBS and resuspended in 15% BSA for 10 minutes. The cells were centrifuged and the cell pellet fixed by slowly adding the fixative for 48 hours at 4°C. The cell pellets were cut in smaller pieces and prepared for EM like following. Pellet fragments were washed in phosphate buffer, stained with 1% osmium tetraoxide in phosphate buffer for 90 minutes, dehydrated in ethanol series and acetone and embedded in TAAB 812 Embedding Resin (T030, TAAB). Semithin (1 µm) sections were cut on Leica Ultracut UCT microtome and stained with toluidine blue. Ultrathin (70 nm) sections were collected on uncoated nickel grids (M200-NI, Electron Microscope Sciences). The grids were stained with 3% uranyl acetate for 15 minutes at 60°C and 3% lead citrate (Leica Ultrostain 2) for 6 minutes at RT. The sections were analyzed with JEM-1400 Plus electron microscope, equipped with Quemsa TEM CCD camera and images obtained using Radius software (TEM Imaging Platform software).

### Survival analysis

Normalized gene expression value for HSPD1 was obtained for lung cancer patient samples from GEO (GSE30219) and mRNA z-score values for the TCGA profile (TCGA LUAD, Cell 2018) from cbioportal platform. Survival curves were generated using Kaplan-Meier estimate with the samples categorized into low- and high-HSPD1 groups based on the median of the HSPD1 gene expression value. Log-rank test was conducted to obtain the significance between the two groups in R software.

### Differential gene expression analysis and correlation

Differential gene expression (DGE) analysis was performed between the KHS101-sensitive and KHS101-resistant cell lines identified from the drug sensitivity analysis. Briefly, the top four KHS101-sensitive (NCI-H460, NCI-H1581, LOU-NH91 and A549) and KHS101-resistant (SK-LU-1, NCI-H1563, BEN and NCI-H838) cell lines were chosen for the analysis. Differential expression between the sensitive and resistant groups was performed using CCLE gene expression profile (GSE36133) with RMA log_2_ signal intensity using GEO2R package in GEO with a moderated t-test parameter. 33 up-regulated and 26 down-regulated genes were fileted with a p-value of <0.05 and log_2_ fold change cutoff of 2. In parallel, Pearson’s correlation analysis was performed between 21 cell lines IC_50_ values of KHS101 and 18,569 genes expression from CCLE. With a correlation value of 0.5 and p-value<0.05, 298 positively and 97 negatively correlated genes were filtered. Then, an overlap analysis was performed with the DGEs list and correlation list to identify a stringent gene list of 12 genes. Correlation analysis between HSPD1 gene expression value obtained from GEO (GSE47206) and KHS101 dose response IC_50_ values was performed in GraphPad Prism 8.

### Differential metabolite analysis

Similar to the differential gene expression analysis, a differential metabolite analysis was performed for the top four KHS101-sensitive and KHS101-resistant group cell lines with the information of 225 log_10_ normalized quantitative metabolites levels across 177 CCLE cell lines (33). 7 differential metabolites were identified with a log_10_ fold change of 2 and p-value < 0.05.

### Gene ontology analysis

Gene ontology analysis was performed for the top 20 gene candidates identified from a global CRISPR knockout screen and with treatment using KHS101 drug. Genes were assessed for the enrichment using GO cellular component 2017b in Enrichr (54).

### Fitness gene and cancer gene dependency

HSPD1 fitness was analyzed as described in (28). HSPD1 gene’s cancer dependency was analyzed in a panel of non-small cell lung cancer cell lines (n=52) from PROJECTDRIVE RNAi screen (27). Using RSA gene level metric of threshold -3, the essentiality of the gene in cell lines was determined.

### In vivo experiments

NSG strain (JAX) was used as experimental model to study the tumor growth of HSPD1 knockdown cells (A549 and H1299). For subcutaneous injections, 0.5×10^6^ cells re-suspended in 50 μL 0.9% NaCl were mixed with Matrigel (Corning) in a ratio 1:1 (v:v). Cells were injected in flanks of 9–15-weeks-old NSG with 8 mice per group. Caliper measurements were taken twice a week and tumor volume was calculated using the formula (Length × Width2 × π)/6. After 4/5 weeks, mice were euthanized by cervical dislocation and tumors were isolated and weighted.

C57BL/6 strain was used as experimental model to evaluate KHS101 effect on lung metastasis tumor formation. For tail-vein metastasis assay, 0.5×10^6^ LL2 cells were re-suspended in 100 μl PBS and injected in the tail vein of female C57BL/6, with 10 mice per group. After 2 weeks either vehicle (5% (v/v) Ethanol - 15% (w/v) (2-Hydroxypropyl)-β-cyclo-dextrin (Captisol Technology)) or 6 mg/kg KHS101 (Cellagen Technology) were injected twice a day subcutaneously for 10 days. Lung metastases were monitored by bioluminescence imaging (BLI). Anesthetized mice were intraperitoneally injected with 50 mg/mL D-luciferin (Kayman Chemicals). Bioluminescence images were acquired with Lumina III in vivo Imaging System (IVIS, Perkin Elmer). Time of survival with disease was calculated as difference between the day of death and the day of appearance of the bioluminescence signal.

In vivo experiments were performed by skilled experimenters trained according to FELASA guidelines. Animal protocols were approved by the Institutional Animal Care and Use Committee of the Regierung von Unterfranken.

---

**Supplementary Table 1.**

| **CL-100 ProLiFiler screening** |  |
| --- | --- |
| **Cell line** | **Medium** |
| A549 | DMEM + 10% FCS |
| BEN | DMEM + 10% FCS |
| COR-L279 | RPMI-1640+10% FCS |
| DV90 | RPMI-1640 + 10% FCS |
| EPLC-272H | RPMI-1640+10% FCS |
| H1299 | DMEM + 10% FCS |
| H460 | DMEM + 10% FCS |
| HCC827 | RPMI-1640+10% FCS |
| LOU-NH91 | RPMI-1640+10% FCS |
| NCI-H1048 | RPMI-1640 + 10% FCS |
| NCI-H1437 | RPMI-1640 + 10% FCS |
| NCI-H1563 | RPMI-1640+10% FCS |
| NCI-H1573 | RPMI-1640+10% FCS |
| NCI-H1581 | RPMI-1640 + 10% FCS |
| NCI-H1703 | RPMI-1640 + 10% FCS |
| NCI-H1838 | RPMI-1640+10% FCS |
| NCI-H2009 | RPMI-1640+10% FCS |
| NCI-H2110 | RPMI-1640 + 10% FCS |
| NCI-H2286 | RPMI-1640+10% FCS |
| NCI-H292 | RPMI-1640+10% FCS |
| NCI-H441 | RPMI-1640+10% FCS |
| NCI-H838 | RPMI-1640 + 10% FCS |
| RERF-LC-Ad2 | RPMI-1640+10% FCS |
| RERF-LC-MS | RPMI-1640 + 10% FCS |
| SCLC-21H | DMEM + 10% FCS |
| SK-LU-1 | DMEM + 10% FCS |

**Supplementary Table 2.**

| **HSPD1 sensitivity score from**<br>**PROJECTDRIVE** |  |
| --- | --- |
| **Cell line** | **Sensitivity Score** |
| NCIH1568 | -11.94 |
| NCIH1944 | -11.06 |
| CALU6 | -10.19 |
| NCIH1437 | -10.18 |
| NCIH1299 | -9.45 |
| NCIH2172 | -9.44 |
| NCIH1355 | -8.68 |
| NCIH23 | -8.61 |
| HCC15 | -8.45 |
| NCIH1793 | -8.29 |
| KNS62 | -7.59 |
| HCC1359 | -6.82 |
| NCIH441 | -6.63 |
| NCIH358 | -6.52 |
| EBC1 | -6.44 |
| NCIH1703 | -6.15 |
| ABC1 | -6.10 |
| NCIH2030 | -6.03 |
| CORL23 | -5.76 |
| RERFLCMS | -5.65 |
| HCC4006 | -5.25 |
| NCIH2110 | -5.18 |
| A549 | -4.47 |
| SQ1 | -4.40 |
| NCIH522 | -4.20 |
| HCC44 | -4.17 |
| NCIH2170 | -4.16 |
| NCIH1693 | -4.02 |
| NCIH661 | -4.01 |
| NCIH1792 | -3.97 |
| NCIH1373 | -3.74 |
| NCIH460 | -3.66 |
| NCIH2122 | -3.64 |
| NCIH2009 | -3.63 |
| LCLC103H | -3.63 |
| SW1573 | -3.43 |
| NCIH1975 | -3.23 |

**Supplementary Table 3.**

| **Mutation profile of the top 4 more sensitive and resistant cell lines** |  |  |  |  |  |  |  |  |
| --- | --- | --- | --- | --- | --- | --- | --- | --- |
| **Gene** | **NCI-H460** | **NCI-H1581** | **LOU-H91** | **A549** | **SK-LU-1** | **NCI-H1563** | **BEN** | **NCI-H838** |
| TP53 | NM | NS<br>(p.Q144*) | MS<br>(p.V143M) | NM | MS<br>(p.H193R) | NM | MS<br>(p.Y163C) | NS<br>(p.E62*) |
| PIK3CA | MS<br>(p.E545K) | NM | MS<br>(p.E726K) | NM | NM | MS<br>(p.E542K) | MS<br>(p.G967R) | NM |
| KRAS | MS<br>(p.Q61H) | NM | NM | MS<br>(p.G12S) | MS<br>(p.G12D) | NM | NM | NM |
| STK11 | NS<br>(p.Q37*) | NM | NM | NS<br>(p.Q37*) | NM | MS<br>(p.G242W),<br>NS (p.Y272*) | NM | NM |

NM = no mutation
MS = missense
NS = nonsense

**Supplementary Table 4.**

| **4-Fold up-regulated genes in resistant cell lines** |  |  |
| --- | --- | --- |
| **Genes** | **logFC** | **p-value** |
| CPE | 5.01 | 0.0033 |
| ACKR3 | 4.13 | 0.0073 |
| GRAMD3 | 3.97 | 0.0003 |
| HCP5 | 3.74 | 0.0289 |
| FOS | 3.47 | 0.0257 |
| NDRG1 | 3.04 | 0.0053 |
| CARD16 | 2.96 | 0.0271 |
| HLA-F | 2.93 | 0.0332 |
| PSMB8-AS1 | 2.82 | 0.0346 |
| ARMCX1 | 2.82 | 0.0389 |
| SATB1 | 2.70 | 0.0231 |
| LNX1 | 2.68 | 0.0350 |
| TMTC1 | 2.60 | 0.0075 |
| MCOLN3 | 2.59 | 0.0035 |
| PLPP3 | 2.57 | 0.0163 |
| TNFRSF21 | 2.57 | 0.0219 |
| PPARGC1A | 2.53 | 0.0493 |
| MSC-AS1 | 2.48 | 0.0466 |
| NLRC5 | 2.43 | 0.0052 |
| TMCC3 | 2.41 | 0.0083 |
| TNFSF10 | 2.37 | 0.0427 |
| UST | 2.36 | 0.0109 |
| DNAJA4 | 2.35 | 0.0491 |
| NMNAT2 | 2.31 | 0.0233 |
| LINC00648 | 2.25 | 0.0112 |
| HLA-J | 2.24 | 0.0250 |
| TAPBPL | 2.24 | 0.0237 |
| TLE2 | 2.21 | 0.0394 |
| ST3GAL6 | 2.14 | 0.0340 |
| AK4 | 2.12 | 0.0500 |
| PRSS21 | 2.08 | 0.0352 |
| HLA-DMA | 2.07 | 0.0487 |
| NDNF | 2.05 | 0.0324 |

**Supplementary Table 5.**

| **4-Fold down-regulated genes in resistant cell lines** |  |  |
| --- | --- | --- |
| **Genes** | **logFC** | **p-value** |
| SH3KBP1 | -2.11 | 0.0186 |
| C11ORF70 | -2.13 | 0.0329 |
| TGFB2 | -2.15 | 0.0373 |
| ECHDC3 | -2.17 | 0.0144 |
| SLFN11 | -2.22 | 0.0443 |
| HHEX | -2.22 | 0.0003 |
| MTAP | -2.23 | 0.0127 |
| GSAP | -2.24 | 0.0304 |
| FOXF2 | -2.31 | 0.0195 |
| SNHG5 | -2.37 | 0.0004 |
| HIST1H1D | -2.39 | 0.0063 |
| FAM133A | -2.48 | 0.0470 |
| RFLNB | -2.49 | 0.0144 |
| NRXN3 | -2.53 | 0.0223 |
| CEP112 | -2.53 | 0.0404 |
| SMKR1 | -2.55 | 0.0057 |
| P2RX5 | -2.57 | 0.0083 |
| SLC6A8 | -2.61 | 0.0151 |
|  | -2.67 | 0.0055 |
| SEMA3A | -2.84 | 0.0080 |
| GLB1L2 | -3.02 | 0.0031 |
| EEF1A2 | -3.23 | 0.0356 |
| LDOC1 | -3.57 | 0.0231 |
| CDH11 | -4.39 | 0.0208 |
| VIM | -4.56 | 0.0477 |
| COL5A2 | -5.54 | 0.0120 |
| GJA1 | -7.11 | 0.0001 |

**Supplementary Table 6.**
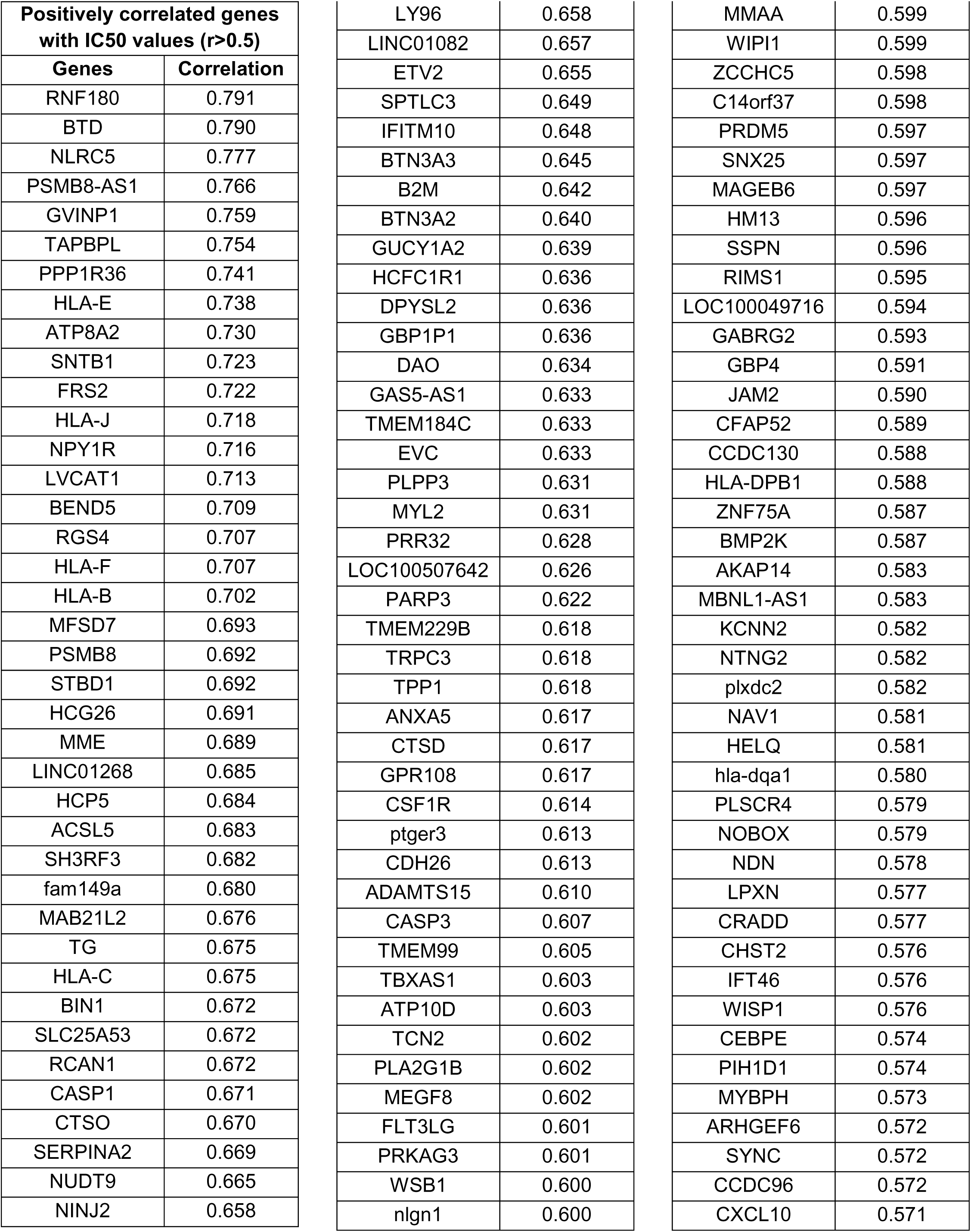
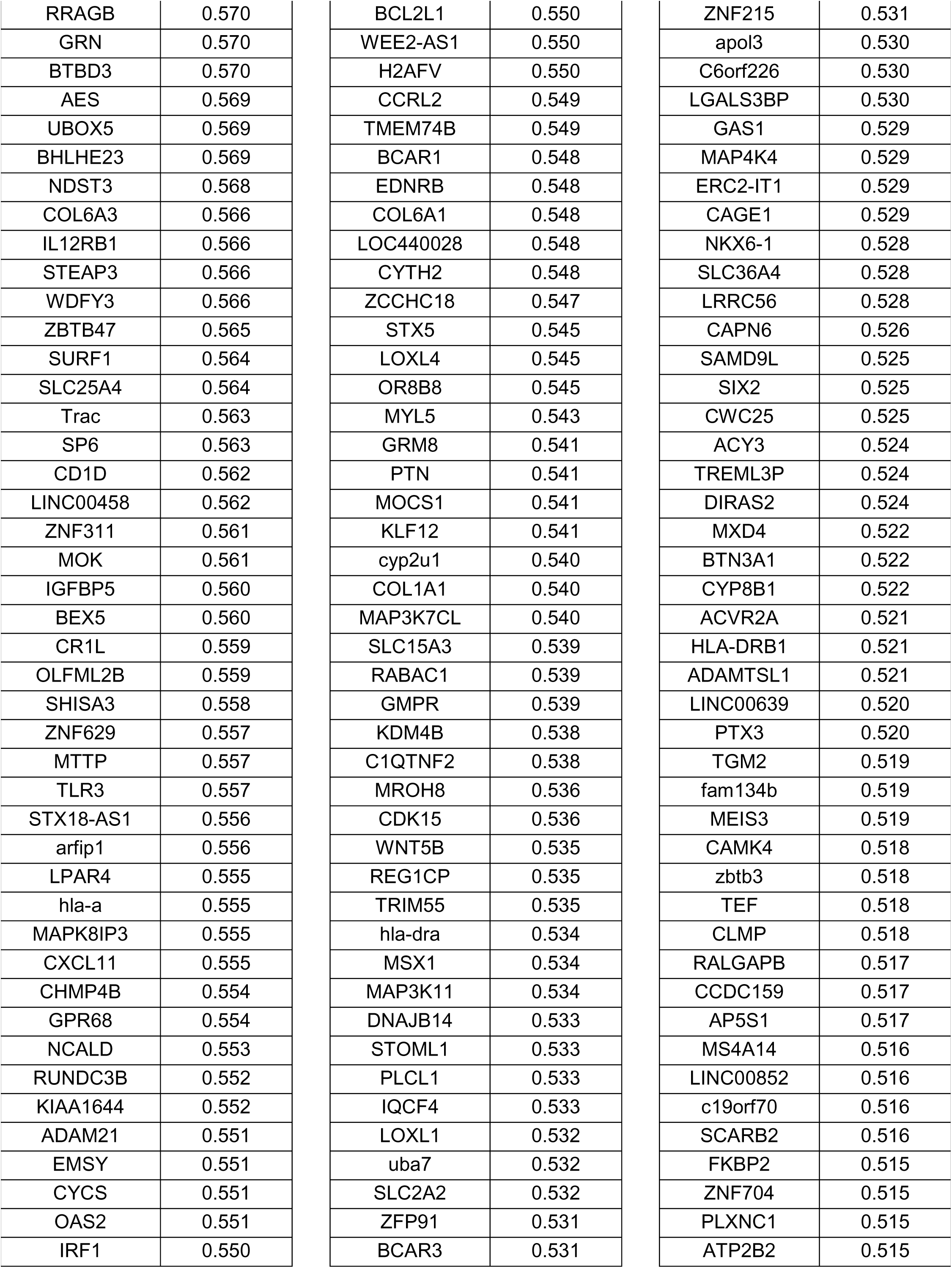
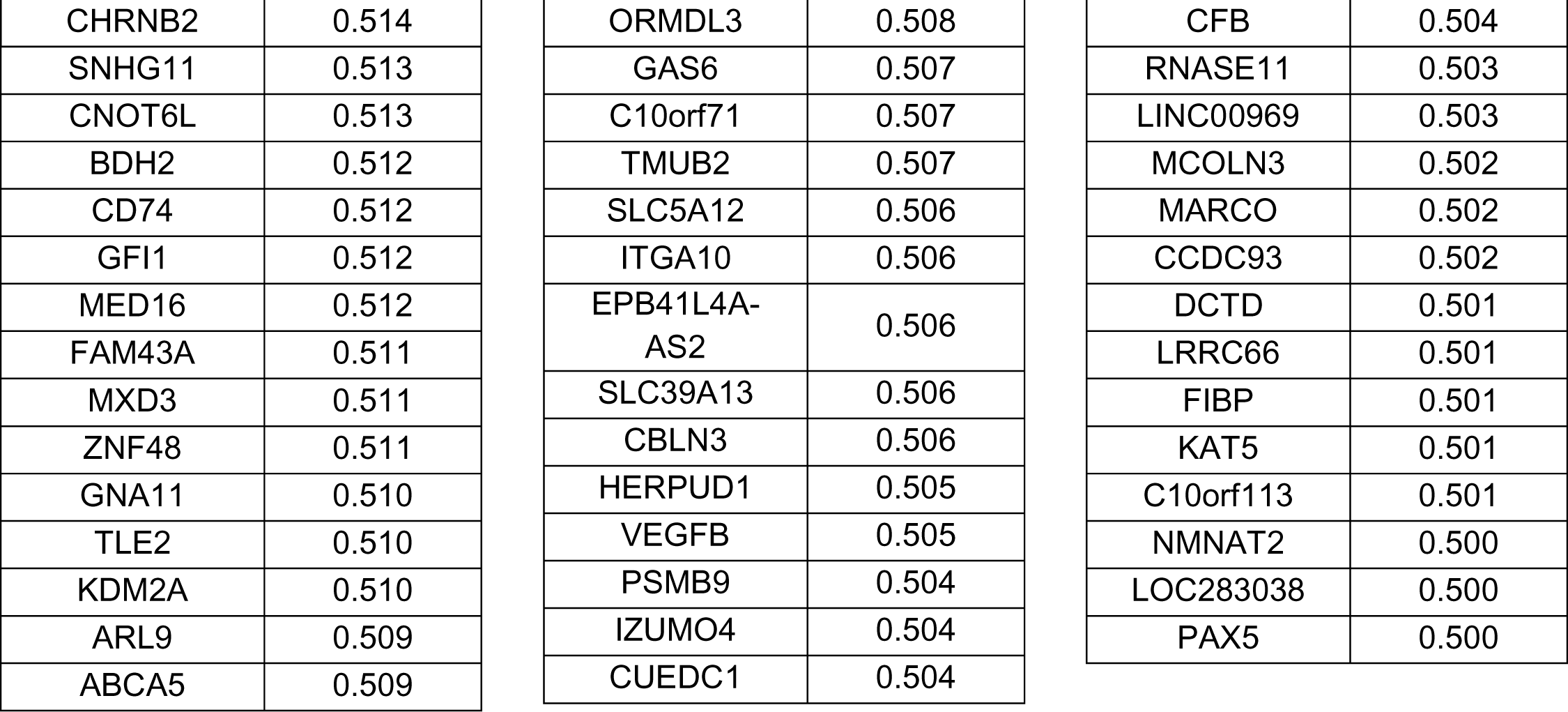

**Supplementary Table 7.**

| Negatively correlated genes with IC50 values ($r < -0.5$) | |
| --- | --- |
| Genes | Correlation |
| ANKS6 | -0.746 |
| GLRX3 | -0.741 |
| RPL13 | -0.698 |
| FAM206A | -0.683 |
| LYNX1 | -0.672 |
| WBSCR22 | -0.662 |
| LOC440704 | -0.660 |
| CTU2 | -0.659 |
| c10orf76 | -0.657 |
| DUS4L | -0.641 |
| PPA1 | -0.627 |
| FN3K | -0.623 |
| POLA1 | -0.621 |
| EIF3A | -0.615 |
| MRPL43 | -0.615 |
| POLR2A | -0.608 |
| SLC6A8 | -0.604 |
| NKD1 | -0.603 |
| EIF1AX | -0.602 |
| armc9 | -0.598 |
| ZCCHC9 | -0.597 |
| IKBKAP | -0.593 |
| RXFP1 | -0.589 |
| PUDP | -0.589 |
| GBF1 | -0.583 |
| NOLC1 | -0.582 |
| SHH | -0.575 |
| ADPGK | -0.575 |
| DUSP22 | -0.575 |
| HAND1 | -0.571 |
| PGAM5 | -0.571 |
| hyal3 | -0.565 |
| HTR1A | -0.564 |
| LINC00544 | -0.564 |
| SLC15A4 | -0.563 |
| LDLRAP1 | -0.563 |
| ANKLE1 | -0.563 |
| RRP9 | -0.562 |
| aire | -0.559 |
| pelp1 | -0.558 |
| COG5 | -0.555 |
| zfp64 | -0.554 |
| RRP12 | -0.553 |
| ACP1 | -0.553 |
| RHBDF2 | -0.552 |
| RIT2 | -0.551 |
| ARMC10 | -0.549 |
| MRGPRG-AS1 | -0.545 |
| SMYD5 | -0.544 |
| EIF2S3 | -0.542 |
| SIGMAR1 | -0.541 |
| USP54 | -0.540 |
| ALDH18A1 | -0.539 |
| NUP93 | -0.539 |
| PDC | -0.539 |
| DHODH | -0.537 |
| HPS6 | -0.536 |
| TCOF1 | -0.536 |
| TNFRSF4 | -0.535 |
| BCCIP | -0.534 |
| TRAV12-1 | -0.533 |
| PES1 | -0.533 |
| HSD17B3 | -0.532 |
| PAPOLB | -0.532 |
| dhx37 | -0.529 |
| NEUROG3 | -0.529 |
| TFB1M | -0.529 |
| SMC1A | -0.527 |
| NME1 | -0.527 |
| RSG1 | -0.527 |
| BLOC1S2 | -0.521 |
| P2RX5 | -0.521 |
| C1QBP | -0.519 |
| ODF1 | -0.518 |
| ttll13p | -0.518 |
| BTNL2 | -0.517 |
| NUP88 | -0.516 |
| MRM1 | -0.515 |
| IQCK | -0.515 |
| pprc1 | -0.514 |
| CD163 | -0.512 |
| IARS | -0.511 |
| rpp25 | -0.510 |
| DDX51 | -0.510 |
| OGFOD1 | -0.510 |
| SLC25A13 | -0.508 |
| ARID1A | -0.507 |
| ZDHHC16 | -0.507 |
| C12ORF10 | -0.507 |
| INPP5A | -0.507 |
| NOC4L | -0.507 |
| LOC100506405 | -0.506 |
| RPL38 | -0.504 |
| CHRNA1 | -0.504 |
| DIMT1 | -0.504 |
| EXOSC1 | -0.503 |
| PDCD11 | -0.501 |

**Supplementary Table 8.**

| **Top 20 genes in CRISPR/Cas9 screening** |  |
| --- | --- |
| **Gene** | **Pos-score** |
| COX5B | 0.000016396 |
| MARK4 | 0.000025227 |
| TATDN3 | 0.000030289 |
| CTAGE1 | 0.000040364 |
| CBR1 | 0.000044941 |
| hsa-mir-365a | 0.000051036 |
| TMC8 | 0.000056552 |
| hsa-mir-138-1 | 0.000066093 |
| SLC25A52 | 0.000070277 |
| CCDC106 | 0.000075681 |
| STK39 | 0.000077296 |
| hsa-mir-181d | 0.000096834 |
| hsa-mir-6856 | 0.00011253 |
| hsa-mir-7850 | 0.00011772 |
| LIPG | 0.00012613 |
| hsa-mir-6730 | 0.00015096 |
| hsa-mir-1184-3 | 0.00015597 |
| hsa-mir-548f-5 | 0.00018152 |
| ACSL5 | 0.00020533 |
| PDE2A | 0.00021423 |

## REFERENCES

1. Bray F, Ferlay J, Soerjomataram I, Siegel RL, Torre LA, Jemal A. Global cancer statistics 2018: GLOBOCAN estimates of incidence and mortality worldwide for 36 cancers in 185 countries. CA Cancer J Clin 2018;68:394–424

2. Herbst RS, Morgensztern D, Boshoff C. The biology and management of non-small cell lung cancer. Nature 2018;553:446–54

3. Kleczko EK, Kwak JW, Schenk EL, Nemenoff RA. Targeting the Complement Pathway as a Therapeutic Strategy in Lung Cancer. Front Immunol 2019;10

4. Osmani L, Askin F, Gabrielson E, Li QK. Current WHO guidelines and the critical role of immunohistochemical markers in the subclassification of non-small cell lung carcinoma (NSCLC): Moving from targeted therapy to immunotherapy. Semin Cancer Biol 2018;52:103–9

5. Duma N, Santana-Davila R, Molina JR. Non-Small Cell Lung Cancer: Epidemiology, Screening, Diagnosis, and Treatment. Mayo Clin Proc 2019;94:1623–40

6. Luengo A, Abbott KL, Davidson SM, Hosios AM, Faubert B, Chan SH, et al. Reactive metabolite production is a targetable liability of glycolytic metabolism in lung cancer. Nat Commun 2019;10:5604

7. Vander Heiden MG, DeBerardinis RJ. Understanding the Intersections between Metabolism and Cancer Biology. Cell 2017;168:657–69

8. Yu L, Lu M, Jia D, Ma J, Ben-Jacob E, Levine H, et al. Modeling the Genetic Regulation of Cancer Metabolism: Interplay between Glycolysis and Oxidative Phosphorylation. Cancer Res 2017;77:1564–74

9. Faubert B, Solmonson A, DeBerardinis RJ. Metabolic reprogramming and cancer progression. Science 2020;368

10. Kim J, DeBerardinis RJ. Mechanisms and Implications of Metabolic Heterogeneity in Cancer. Cell Metab 2019;30:434–46

11. Jin N, Bi A, Lan X, Xu J, Wang X, Liu Y, et al. Identification of metabolic vulnerabilities of receptor tyrosine kinases-driven cancer. Nat Commun 2019;10:2701

12. Fenouille N, Bassil CF, Ben-Sahra I, Benajiba L, Alexe G, Ramos A, et al. The creatine kinase pathway is a metabolic vulnerability in EVI1-positive acute myeloid leukemia. Nat Med 2017;23:301–13

13. Bader DA, Hartig SM, Putluri V, Foley C, Hamilton MP, Smith EA, et al. Mitochondrial pyruvate import is a metabolic vulnerability in androgen receptor-driven prostate cancer. Nat Metab 2019;1:70–85

14. Jacob P, Hirt H, Bendahmane A. The heat-shock protein/chaperone network and multiple stress resistance. Plant Biotechnol J 2017;15:405–14

15. Dubrez L, Causse S, Borges Bonan N, Dumetier B, Garrido C. Heat-shock proteins: chaperoning DNA repair. Oncogene 2020;39:516–29

16. Chatterjee S, Burns TF. Targeting Heat Shock Proteins in Cancer: A Promising Therapeutic Approach. Int J Mol Sci 2017;18

17. Wu J, Liu T, Rios Z, Mei Q, Lin X, Cao S. Heat Shock Proteins and Cancer. Trends Pharmacol Sci 2017;38:226–56

18. Gomez-Llorente Y, Jebara F, Patra M, Malik R, Nisemblat S, Chomsky-Hecht O, et al. Structural basis for active single and double ring complexes in human mitochondrial Hsp60-Hsp10 chaperonin. Nat Commun 2020;11:1916

19. Teng R, Liu Z, Tang H, Zhang W, Chen Y, Xu R, et al. HSP60 silencing promotes Warburg-like phenotypes and switches the mitochondrial function from ATP production to biosynthesis in ccRCC cells. Redox Biol 2019;24:101218-

20. Yun CW, Kim HJ, Lim JH, Lee SH. Heat Shock Proteins: Agents of Cancer Development and Therapeutic Targets in Anti-Cancer Therapy. Cells 2019;9

21. Xu X, Wang W, Shao W, Yin W, Chen H, Qiu Y, et al. Heat shock protein-60 expression was significantly correlated with the prognosis of lung adenocarcinoma. J Surg Oncol 2011;104:598–603

22. Fucarino A, Pitruzzella A. Role of HSP60/HSP10 in Lung Cancer: Simple Biomarkers or Leading Actors? J Oncol 2020;2020:4701868

23. Polson ES, Kuchler VB, Abbosh C, Ross EM, Mathew RK, Beard HA, et al. KHS101 disrupts energy metabolism in human glioblastoma cells and reduces tumor growth in mice. Sci Transl Med 2018;10

24. Siddiqui A, Gollavilli PN, Schwab A, Vazakidou ME, Ersan PG, Ramakrishnan M, et al. Thymidylate synthase maintains the de-differentiated state of triple negative breast cancers. Cell Death Differ 2019;26:2223–36

25. Gollavilli PN, Parma B, Siddiqui A, Yang H, Ramesh V, Napoli F, et al. The role of miR-200b/c in balancing EMT and proliferation revealed by an activity reporter. Oncogene 2021

26. Behan FM, Iorio F, Picco G, Goncalves E, Beaver CM, Migliardi G, et al. Prioritization of cancer therapeutic targets using CRISPR-Cas9 screens. Nature 2019;568:511–6

27. McDonald ER, 3rd, de Weck A, Schlabach MR, Billy E, Mavrakis KJ, Hoffman GR, et al. Project DRIVE: A Compendium of Cancer Dependencies and Synthetic Lethal Relationships Uncovered by Large-Scale, Deep RNAi Screening. Cell 2017;170:577–92 e10

28. Siddiqui MA, Gollavilli PN, Ramesh V, Parma B, Schwab A, Vazakidou ME, et al. Thymidylate synthase drives the phenotypes of epithelial-to-mesenchymal transition in non-small cell lung cancer. Br J Cancer 2021;124:281–9

29. Senan S, Brade A, Wang LH, Vansteenkiste J, Dakhil S, Biesma B, et al. PROCLAIM: Randomized Phase III Trial of Pemetrexed-Cisplatin or Etoposide-Cisplatin Plus Thoracic Radiation Therapy Followed by Consolidation Chemotherapy in Locally Advanced Nonsquamous Non-Small-Cell Lung Cancer. J Clin Oncol 2016;34:953–62

30. Kim SJ, Kim HS, Seo YR. Understanding of ROS-Inducing Strategy in Anticancer Therapy. Oxid Med Cell Longev 2019;2019:5381692

31. Benko S, Kovacs EG, Hezel F, Kufer TA. NLRC5 Functions beyond MHC I Regulation-What Do We Know So Far? Front Immunol 2017;8:150

32. Ji L, Zhao X, Zhang B, Kang L, Song W, Zhao B, et al. Slc6a8-Mediated Creatine Uptake and Accumulation Reprogram Macrophage Polarization via Regulating Cytokine Responses. Immunity 2019;51:272–84 e7

33. Li H, Ning S, Ghandi M, Kryukov GV, Gopal S, Deik A, et al. The landscape of cancer cell line metabolism. Nat Med 2019;25:850–60

34. Snow RJ, Murphy RM. Creatine and the creatine transporter: a review. Mol Cell Biochem 2001;224:169–81

35. Ramesh V, Brabletz T, Ceppi P. Targeting EMT in Cancer with Repurposed Metabolic Inhibitors. Trends in cancer 2020;6:942–50

36. Kawada K, Toda K, Sakai Y. Targeting metabolic reprogramming in KRAS-driven cancers. Int J Clin Oncol 2017;22:651–9

37. Obrist F, Michels J, Durand S, Chery A, Pol J, Levesque S, et al. Metabolic vulnerability of cisplatin-resistant cancers. The EMBO journal 2018;37

38. Marino Gammazza A, Macaluso F, Di Felice V, Cappello F, Barone R. Hsp60 in Skeletal Muscle Fiber Biogenesis and Homeostasis: From Physical Exercise to Skeletal Muscle Pathology. Cells 2018;7

39. Tsai YP, Yang MH, Huang CH, Chang SY, Chen PM, Liu CJ, et al. Interaction between HSP60 and beta-catenin promotes metastasis. Carcinogenesis 2009;30:1049–57

40. Berger E, Rath E, Yuan D, Waldschmitt N, Khaloian S, Allgäuer M, et al. Mitochondrial function controls intestinal epithelial stemness and proliferation. Nat Commun 2016;7:13171

41. Michils A, Redivo M, Zegers de Beyl V, de Maertelaer V, Jacobovitz D, Rocmans P, et al. Increased expression of high but not low molecular weight heat shock proteins in resectable lung carcinoma. Lung Cancer 2001;33:59–67

42. Sotgia F, Lisanti MP. Mitochondrial markers predict survival and progression in non-small cell lung cancer (NSCLC) patients: Use as companion diagnostics. Oncotarget 2017;8:68095–107

43. Zhou C, Sun H, Zheng C, Gao J, Fu Q, Hu N, et al. Oncogenic HSP60 regulates mitochondrial oxidative phosphorylation to support Erk1/2 activation during pancreatic cancer cell growth. Cell Death Dis 2018;9:161

44. Wurdak H, Zhu S, Min KH, Aimone L, Lairson LL, Watson J, et al. A small molecule accelerates neuronal differentiation in the adult rat. Proc Natl Acad Sci U S A 2010;107:16542–7

45. Lim ZF, Ma PC. Emerging insights of tumor heterogeneity and drug resistance mechanisms in lung cancer targeted therapy. J Hematol Oncol 2019;12:134

46. Pirker R. Chemotherapy remains a cornerstone in the treatment of nonsmall cell lung cancer. Curr Opin Oncol 2020;32:63–7

47. Visconti R, Morra F, Guggino G, Celetti A. The between Now and Then of Lung Cancer Chemotherapy and Immunotherapy. Int J Mol Sci 2017;18

48. Scagliotti GV, Ceppi P, Capelletto E, Novello S. Updated clinical information on multitargeted antifolates in lung cancer. Clin Lung Cancer 2009;10 Suppl 1:S35–40

49. Khan S, He Y, Zhang X, Yuan Y, Pu S, Kong Q, et al. PROteolysis TArgeting Chimeras (PROTACs) as emerging anticancer therapeutics. Oncogene 2020;39:4909–24

50. Sun X, Gao H, Yang Y, He M, Wu Y, Song Y, et al. PROTACs: great opportunities for academia and industry. Signal Transduct Target Ther 2019;4:64

51. Siddiqui A, Vazakidou ME, Schwab A, Napoli F, Fernandez-Molina C, Rapa I, et al. Thymidylate synthase is functionally associated with ZEB1 and contributes to the epithelial-to-mesenchymal transition of cancer cells. J Pathol 2017;242:221–33

52. Sanjana NE, Shalem O, Zhang F. Improved vectors and genome-wide libraries for CRISPR screening. Nature methods 2014;11:783–4

53. Li W, Xu H, Xiao T, Cong L, Love MI, Zhang F, et al. MAGeCK enables robust identification of essential genes from genome-scale CRISPR/Cas9 knockout screens. Genome Biol 2014;15:554

54. Chen EY, Tan CM, Kou Y, Duan Q, Wang Z, Meirelles GV, et al. Enrichr: interactive and collaborative HTML5 gene list enrichment analysis tool. BMC Bioinformatics 2013;14:128

